# The Orphan G Protein-Coupled Receptor GPR52 is a Novel Regulator of Breast Cancer Multicellular Organization

**DOI:** 10.1101/2024.07.22.604482

**Authors:** Sarah Z. Hanif, CheukMan Cherie Au, Ingrid Torregroza, Caleb Kutz, Syeda Y. Jannath, Tabassum Fabiha, Bhavneet Bhinder, Michael P. Washburn, Dominic Devost, Shuchen Liu, Priya Bhardwaj, Todd Evans, Pradeep K. Anand, Robert Tarran, Sailesh Palikhe, Olivier Elemento, Lukas E. Dow, John Blenis, Terence E. Hébert, Kristy A. Brown

**Author notes:** The authors declare no potential conflicts of interest.

## Abstract

**Statement of Significance:** We showed that loss of the orphan G protein-coupled receptor GPR52 in human breast cell lines leads to increased cell clustering, hybrid/partial EMT, and increased tumor burden in zebrafish.

**Background:** G protein-coupled receptors (GPCRs) are the largest class of membrane-bound receptors that transmit critical signals from extracellular to intracellular spaces. Transcriptomic data of resected breast tumors show that low mRNA expression of orphan GPCR GPR52 correlates with reduced overall survival in patients with breast cancer, leading to the hypothesis that loss of GPR52 supports breast cancer progression.

**Methods:** CRISPR-Cas9 was used to knockout GPR52 in the human triple-negative breast cancer (TNBC) cell lines MDA-MB-468 and MDA-MB-231, and in the non-cancerous breast epithelial cell line MCF10A. 2D and 3D *in vitro* studies, electron microscopy, Matrigel culture, and a zebrafish xenograft model were used to assess the morphology and behavior of GPR52 KO cells. RNA-sequencing and proteomic analyses were also conducted on these cell lines, and transcriptomic data from The Cancer Genome Atlas (TCGA) database were used to compare GPR52-null and wild-type (WT) signatures in breast cancer.

**Results:** Loss of GPR52 was found to be associated with increased cell-cell interaction in 2D cultures, altered 3D spheroid morphology, and increased propensity to organize and invade collectively in Matrigel. Furthermore, GPR52 loss was associated with features of EMT in MDA-MB-468 cells, and zebrafish injected with GPR52 KO cells developed a greater total cancer area than those injected with control cells. RNA sequencing and proteomic analyses of GPR52-null breast cancer cells revealed an increased cAMP signaling signature. Consistently, we found that treatment of wild-type (WT) cells with forskolin, which stimulates the production of cAMP, induces phenotypic changes associated with GPR52 loss, and inhibition of cAMP production rescued some GPR52 KO phenotypes.

**Conclusion:** GPR52 is an orphan GPCR and its role in cancer progression has not been previously characterized. We found that GPR52 loss in breast cancer cells can lead to increased cell clustering, collective invasion, and EMT *in vitro*. These are features of increased cancer aggression. Our results reveal that GPR52 loss is a potential mechanism by which breast cancer progression may occur and support the investigation of GPR52 agonism as a therapeutic option for breast cancer.

Graphical Abstract

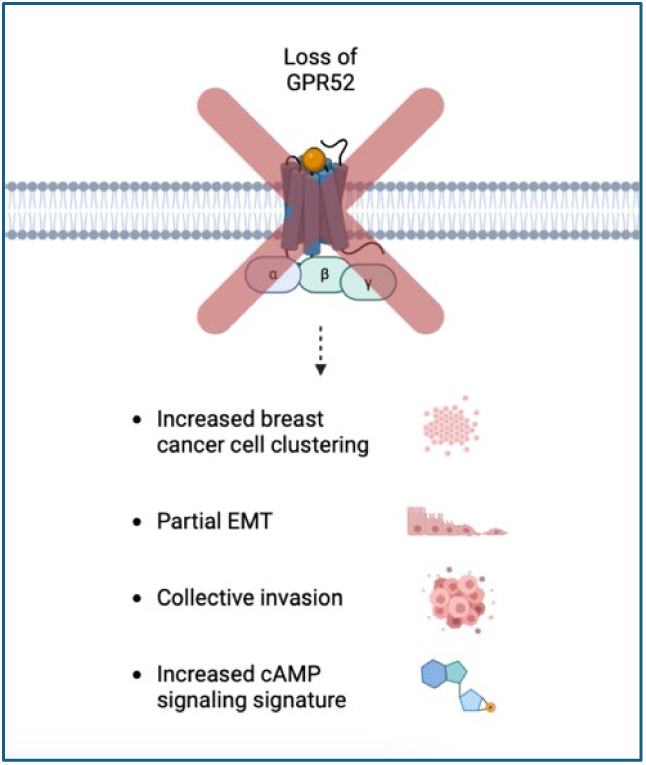

## Background

Metastasis is the primary cause of death in breast cancer patients (1). The process required for cancer cells of solid tumors to metastasize is intensive, and cells undergo several adaptive processes to enhance their metastatic potential. These include changes in cell-cell and cell-matrix adhesion, transitions between epithelial and mesenchymal cell states, and the ability to degrade and invade tissues (2, 3). However, the upstream regulators of these processes are not well characterized, which limits mechanistic understanding and therapeutic intervention.

G protein-coupled receptors (GPCRs) are the largest protein family encoded by the human genome (4). These receptors consist of an extracellular N-terminus followed by seven transmembrane α-helices, which are connected by three intracellular and three extracellular loops and a cytoplasmic C-terminal tail (5). Their transmembrane structure enables the transmission of critical signals between extracellular and intracellular spaces. Dissociation of the heterotrimeric G protein upon GPCR activation can regulate a diverse array of downstream molecules, allowing for the regulation of various cell processes, including proliferation, migration, adhesion, and metabolism (6, 7).

GPR52 is a structurally unique GPCR that is enriched in the basal ganglia and its endogenous ligand remains unknown, rendering it an orphan receptor (8). It has garnered increased attention in recent years owing to its potential as a neurotherapeutic target for schizophrenia and Huntington’s disease (9, 10). We examined GPR52 mRNA levels in 19 solid tumor types and determined that GPR52 is significantly downregulated in tumor samples (11). However, the role of GPR52 in cancer progression has not been reported. In patients with breast cancer, we found that GPR52 expression was further reduced in metastases compared with that in the primary tumor (11). Low GPR52 mRNA expression in resected breast tumors is also associated with a reduction in overall survival (12).

We generated GPR52 KO cancerous and non-cancerous breast epithelial cells from the widely used MDA-MB-468, MDA-MB-231, and MCF10A lines. Loss of GPR52 led to an increase in cell-cell interactions in 2D cultures, with the formation of cell clusters in each cell line. Transmission electron microscopy (TEM) of WT and GPR52 KO cells revealed differences in cell-cell adhesion properties, including the length of the cell-cell interface and the proximity of cells along this interface. In 3D Matrigel cultures, GPR52 KO was associated with changes in the organization and morphology of MDA-MB-468 and MDA-MB-231 spheroids. Furthermore, GPR52 loss increased the propensity of breast cancer cells to organize and invade collectively when cultured in Matrigel. Lastly, we found that the culture of GPR52 KO cells on poly-D-lysine led to partial EMT.

RNA-sequencing and proteomic studies of GPR52-null cells demonstrated the upregulation of several pathways implicated in breast cancer, including cAMP signaling (13). Furthermore, we found that phosphorylation of CREB was increased in GPR52 KO cells and that treatment of WT cells with forskolin, which stimulates the production of cAMP, promotes features associated with GPR52 loss, whereas inhibition of cAMP production rescued some of the GPR52 KO phenotypes.

Overall, our results revealed that GPR52 loss is a potential mechanism by which important processes in breast cancer progression, such as EMT and changes in multicellular organization, may occur. These processes have long been implicated in many solid tumors; however, critical and targetable upstream regulators have not been identified. As GPCRs are the targets of more than 1/3 of FDA-approved small-molecule drugs, these data support the investigation of GPR52 agonism as a viable therapeutic approach in breast cancer (14).

## Results

### GPR52 expression is reduced in cancerous tissue compared to normal, and inversely associated with breast cancer prognosis and metastatic potential

To date, there have been no reports on the physiological role of GPR52 in any type of cancer. However, GPR52 mRNA expression levels have been reported in many transcriptome profiles of resected cancerous and noncancerous tissues. We compared GPR52 mRNA expression levels in normal and tumor samples from the tissues from which solid tumors arose (Fig. 1A) using the TNMplot webtool (11). We found that, in the majority of these tissue types, GPR52 mRNA expression levels were lower in tumors than in non-cancerous samples (Fig. 1A, *P*<0.05, indicated with an asterisk). Importantly, low tumor GPR52 mRNA expression was found to be associated with a reduction in overall survival in patients with breast cancer (Fig. 1B), with the impact of low GPR52 on survival being more pronounced for triple-negative breast cancer (TNBC) (Fig. 1C) (12). GPR52 mRNA expression was also lower in metastatic nodes than in the primary resected tumor (Fig. 1D, p=4.45e-17) (11), which is consistent with GPR52 the differentially expressed in several human breast cell lines with varying metastatic potentials (Fig. 1E).

**Figure 1.**
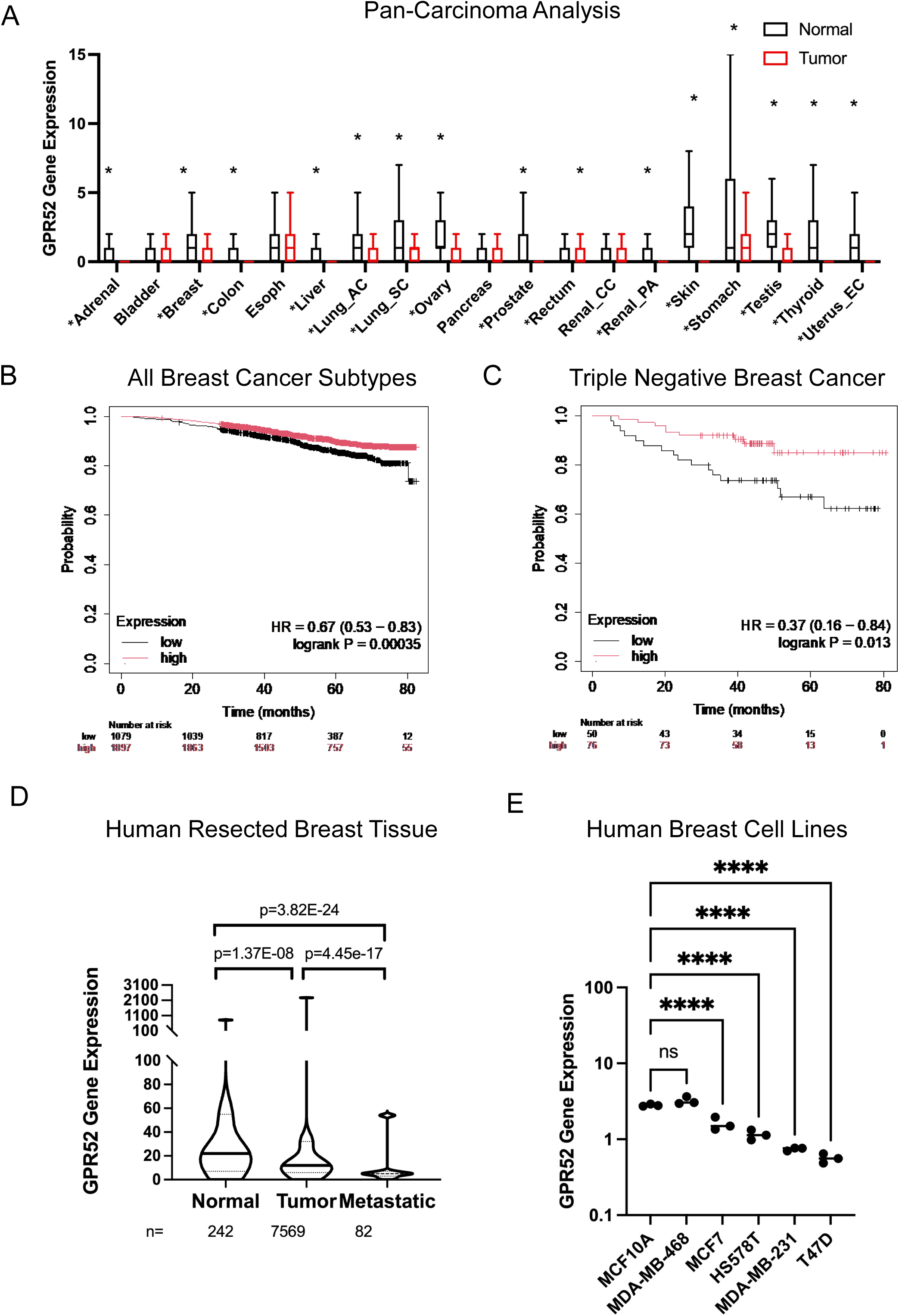
Low GPR52 expression is associated with increased breast cancer progression and reduced survival probability of breast cancer patients. A) The median GPR52 mRNA expression, interquartile range, minimum value, and upper whisker are plotted for normal (black) and tumor (red) tissues. Mann-Whitney U tests were conducted to determine statistical significance. \**P* <0.05. AC=adenocarcinoma, SC=squamous cell, CC=clear cell, PA=papillary cell, EC=endometrial carcinoma. B) KMplot breast cancer overall survival curves for patients with low versus high GPR52 mRNA expression in resected tumors for (B) all breast cancer subtypes and (C) triple negative breast cancer. D) GPR52 mRNA expression collected from a gene chip dataset of non-cancerous breast tissue, primary tumor, and metastases of individuals with breast cancer (un-paired). Data are presented as median with upper and lower quartiles and minimum and maximum values. One-way ANOVA, p<0.05. E) GPR52 mRNA expression was determined by QPCR and normalized to the housekeeping gene RPL32. The normalized GPR52 expression was then divided by the average expression of GPR52 across the cell lines. n=3, line=median. One-way ANOVA, P-value<0.05; \*\*\*\**P* <0.00005, ns=not significant.

### GPR52 regulates cell-cell adhesion properties of cancerous and non-cancerous breast epithelial cells

We used CRISPR-Cas9 to generate indels in *GPR52* in two human breast cancer cell lines (MDA-MB-231 and MDA-MB-468) and one non-cancerous breast epithelial cell line (MCF10A), using two different guide RNAs (Supplementary Fig. S1). The morphology of GPR52 KO cells was notably different from that of control cells. GPR52 KO MDA-MB-231, MDA-MB-468, and MCF10A cell lines generated from both guide RNAs formed clusters of tightly packed cells with clearly defined borders in the monolayer culture (Fig. 2A). To further investigate the effect of GPR52 loss on the interactions between cells, we performed TEM of vector control and GPR52 KO MDA-MB-468 cells cultured in suspension (Fig. 2B). GPR52 KO cells had a shorter cell-cell interface than WT cells (Fig. 2C), but they interacted more closely with one another along the length of their interface than WT cells did (Fig. 2D). The cell diameter was not significantly different between the groups, suggesting that the reduction in cell interface length in the GPR52 KO group was not due to a reduction in cell size (Fig. 2E).

**Figure 2.**
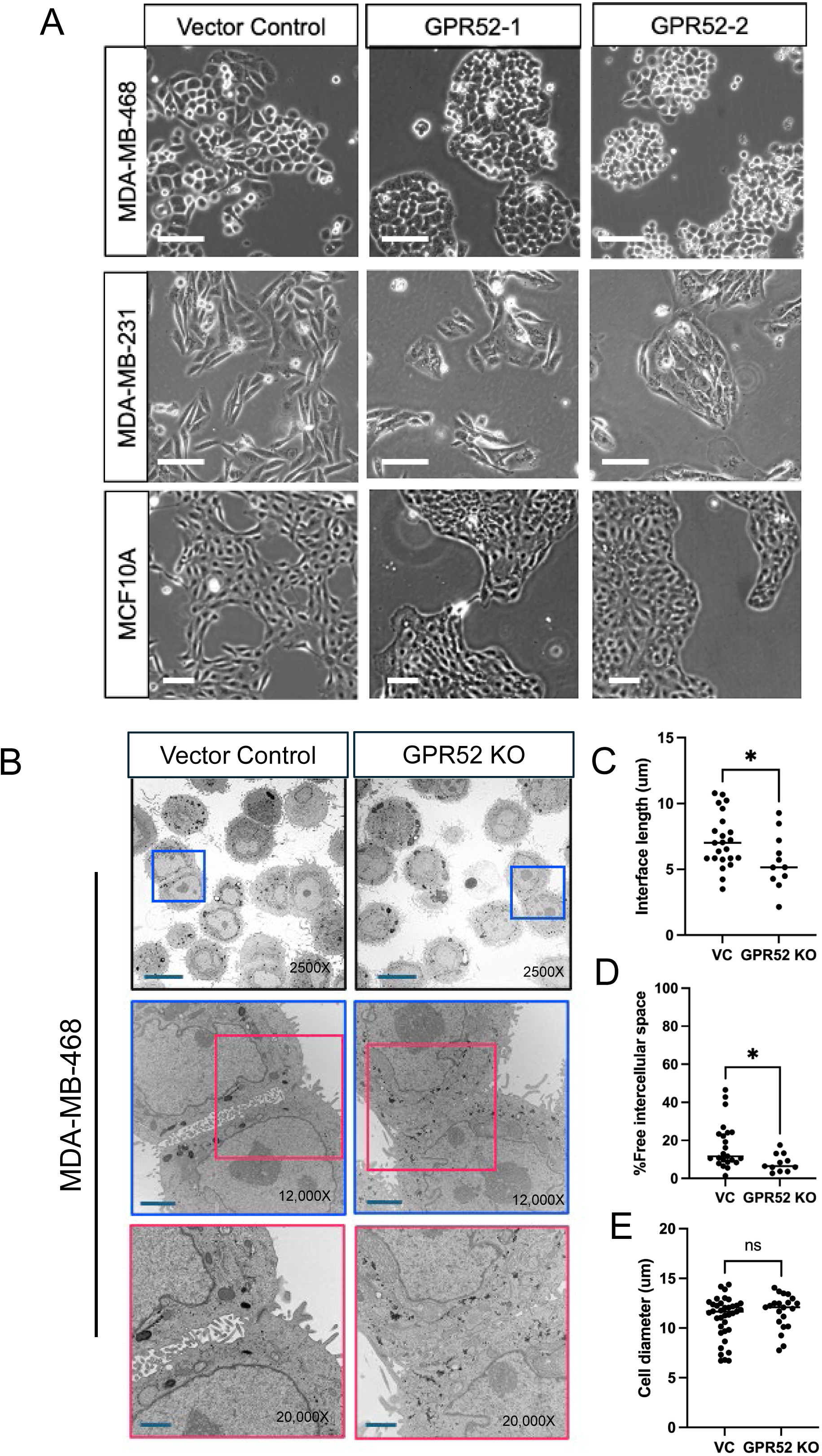
GPR52 regulates cell-cell adhesion properties of cancerous and non-cancerous breast epithelial cells. A) Human breast cell lines were grown on tissue culture-treated plastic under standard cell culture conditions and imaged at 50-60% confluency with light microscopy. Scalebar=100 μm. B) MDA-MB-468 cells cultured in suspension were visualized with TEM from 2500-20,000x magnification. Scalebars: 2500x=10 mm; 12,000x=2 mm, 20,000x=1 mm. C) The length of the interface between cells, D) fraction of the cell-cell interface that was occupied by free space, and E) cell diameter were determined using 2500x images. Student’s t-test, P-value<0.05; \**P* <0.05, ns=not significant. Line=median. VC=vector control.

### GPR52 loss is associated with EMT, collective invasion, and an altered pattern of ECM digestion in breast cancer

Given the rounded property of GPR52 KO cells, we sought to improve their adherence to 2D culture by coating tissue culture plates with poly-D-lysine (Fig. 3A). The culture of GPR52 KO MDA-MB-468 cells on plastic coated with poly-D-lysine led to their elongation and a mesenchymal phenotype. This was not observed in WT cells. Western blotting of GPR52 KO MDA-MB-468 cells revealed upregulation of mesenchymal cell markers, such as Snai1 and vimentin, but the cells continued to express E-cadherin (Fig. 3B).

**Figure 3.**
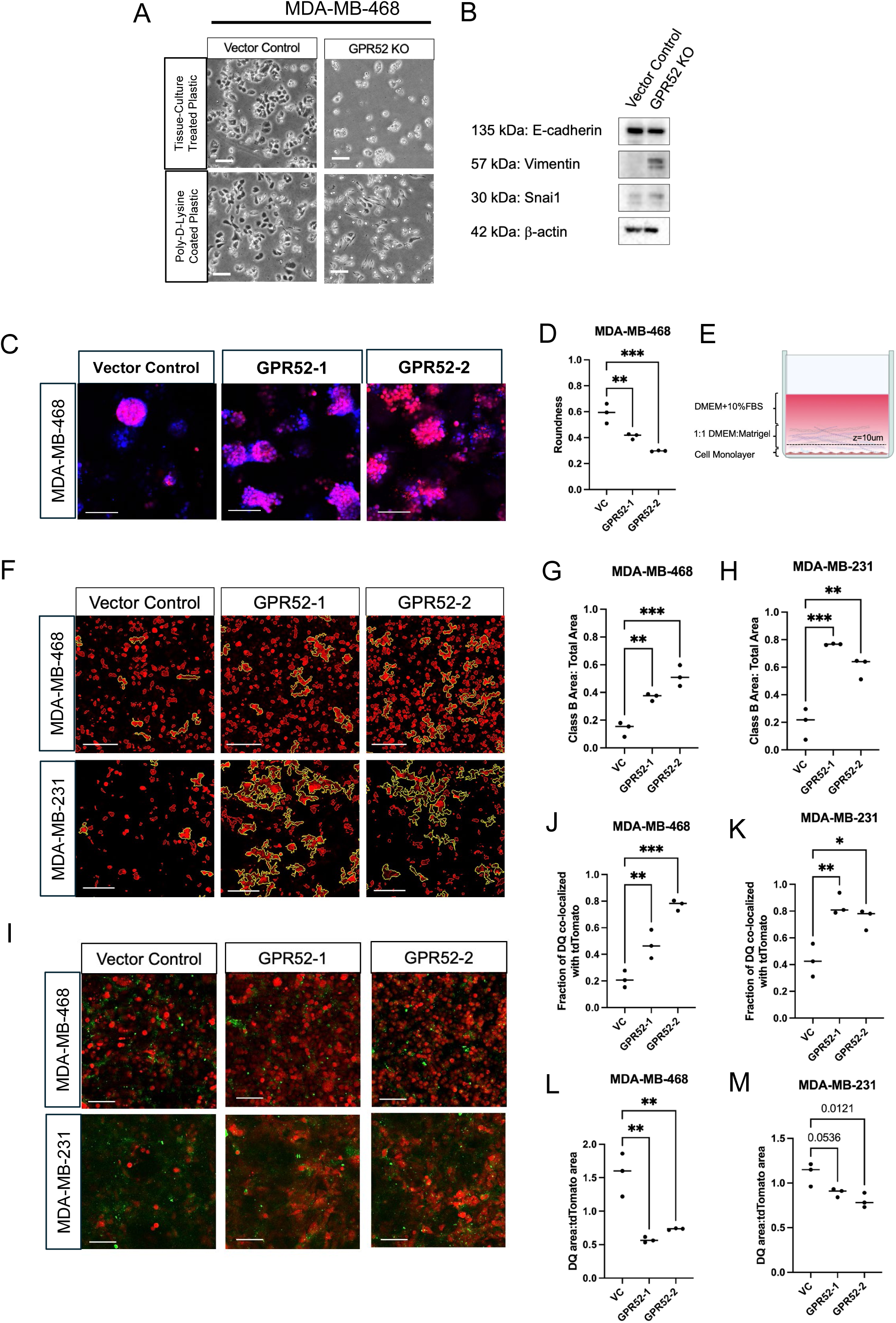
GPR52 loss is associated with EMT, collective invasion, and an altered pattern of ECM digestion in breast cancer. A) Vector control and GPR52 KO (sgRNA2) MDA-MB-468 cells were cultured on tissue-culture treated plates that were untreated (top row) or treated with 1mg/mL poly-D-lysine coated plastic (bottom row). Scalebar=100 mm. B) Western blotting of MDA-MB-468 cells cultured on poly-D-lysine. C) Cells were cultured in Matrigel for 10 days under complete media conditions. Cytoplasmic tdTomato (red) and 1:1000 nuclear Hoechst 33342 stain (blue) allow visualization of MDA-MB-468 WT (vector control) and GPR52 KO (GPR52-1, GPR52-2) spheroids. Scalebar=100 mm. D) The roundness of spheroids was determined based on the tdTomato signal. E) Schematic of invasion assay. FBS=fetal bovine serum, DMEM= Dulbecco’s Modified Eagle Medium. F) Z-stacks obtained at t=24 hours were assessed at z=10 mm. All Class B structures are outlined in yellow. Scalebar=150 μm. The fraction of the area occupied by Class B structures for (G) MDA-MB-468 and (H) MDA-MB-231. VC=vector control. n=3, One-way ANOVA, P-value<0.05. \**P* <0.05, \*\**P* <0.005, ns=not significant. Line=median. I) Representative images are shown of tdTomato-tagged cancer cells (red) and DQ-collagen (green) at t=24 hours at z=10 μm. Scalebar=100 mm. The fraction of DQ co-localized with tdTomato at z=10 mm (J-K) and area of DQ normalized to the area of tdTomato (J-K) were determined for MDA-MB-468 and MDA-MB-231, respectively. n=3, One-way ANOVA, P-value<0.05. \**P* <0.05, \*\**P* <0.005, \*\*\**P* <0.0005, ns=not significant. Line=median.

Compared to WT cells, 3D culture of GPR52 KO MDA-MB-468 cells led to the formation of disorganized spheroids with irregular borders (Fig. 3A-B, top rows; 3C). To further define the impact of GPR52 loss on this invasive phenotype, we plated cells as a monolayer and evaluated their invasion through a layer of Matrigel (Fig. 3E). Interestingly, GPR52 KO cells tended to organize and invade collectively in large clusters (MDA-MB-468) or sheets (MDA-MB-231) (Fig. 3F). We categorized all cancer foci with an area ≥ 3140 μm^2^ at z=10 μm as Class B structures (yellow outline). This threshold was selected because the diameter of MDA-MB-468 and MDA-MB-231 cells was found to range from to 10-20 μm, therefore, the cross-sectional area of approximately 10 cells was 10 × π × r^2^ = 10 × π × 10^2^=∼3140). We then calculated the sum of the areas of all Class B structures and divided this by the sum of the areas of all cancer foci at z=10 μm. We found that the fraction of the area occupied by Class B structures was increased in the MDA-MB-468 and MDA-MB-231 GPR52 KO groups, suggesting an increased propensity to organize collectively (Fig. 3G-H).

Although collective invasion has been investigated previously, the effect of this behavior on the ECM through which cells invade has not been well characterized. To assess the pattern of ECM degradation, we incorporated dye-quenched (DQ)-collagen IV, which contains sequestered fluorophores that are released following proteolysis of collagen IV, a major component of the basement membrane that breast cancer cells invade (18). At the 24-hour timepoint, the DQ signal was found to colocalize strongly with the GPR52 KO cells, while it was more diffuse and less colocalized with the vector control cells for both MDA-MB-468 and MDA-MB-231 cells (Fig. 3I). Furthermore, we normalized the area of the DQ signal to the area occupied by cancer cells and found that it tended to be lower for the GPR52 KO groups than for the vector control, suggesting a focal area of matrix degradation (Fig. 3L-M).

### Proteomic and transcriptomic analyses of GPR52 KO breast cell lines and resected breast tumors

Next, we conducted proteomic analyses of the WT and GPR52 KO MDA-MB-468, MDA-MB-231, and MCF10A cell lines. Using Ingenuity Pathway Analysis (IPA, Qiagen), we observed upregulation of signatures associated with cellular homeostasis, viability, survival, proliferation, and migration in all three cell lines, whereas the signatures associated with organismal death and cell death of tumor cell lines were both significantly reduced (Fig. 4A). Several upstream regulators were predicted to be similarly altered across the GPR52 KO lines (Fig. 4B). We were particularly intrigued by the signature associated with the cAMP analog 8-bromo-cAMP, as GPR52 activation and function have been associated with its regulation of intracellular cAMP levels (9, 10).

**Figure 4.**
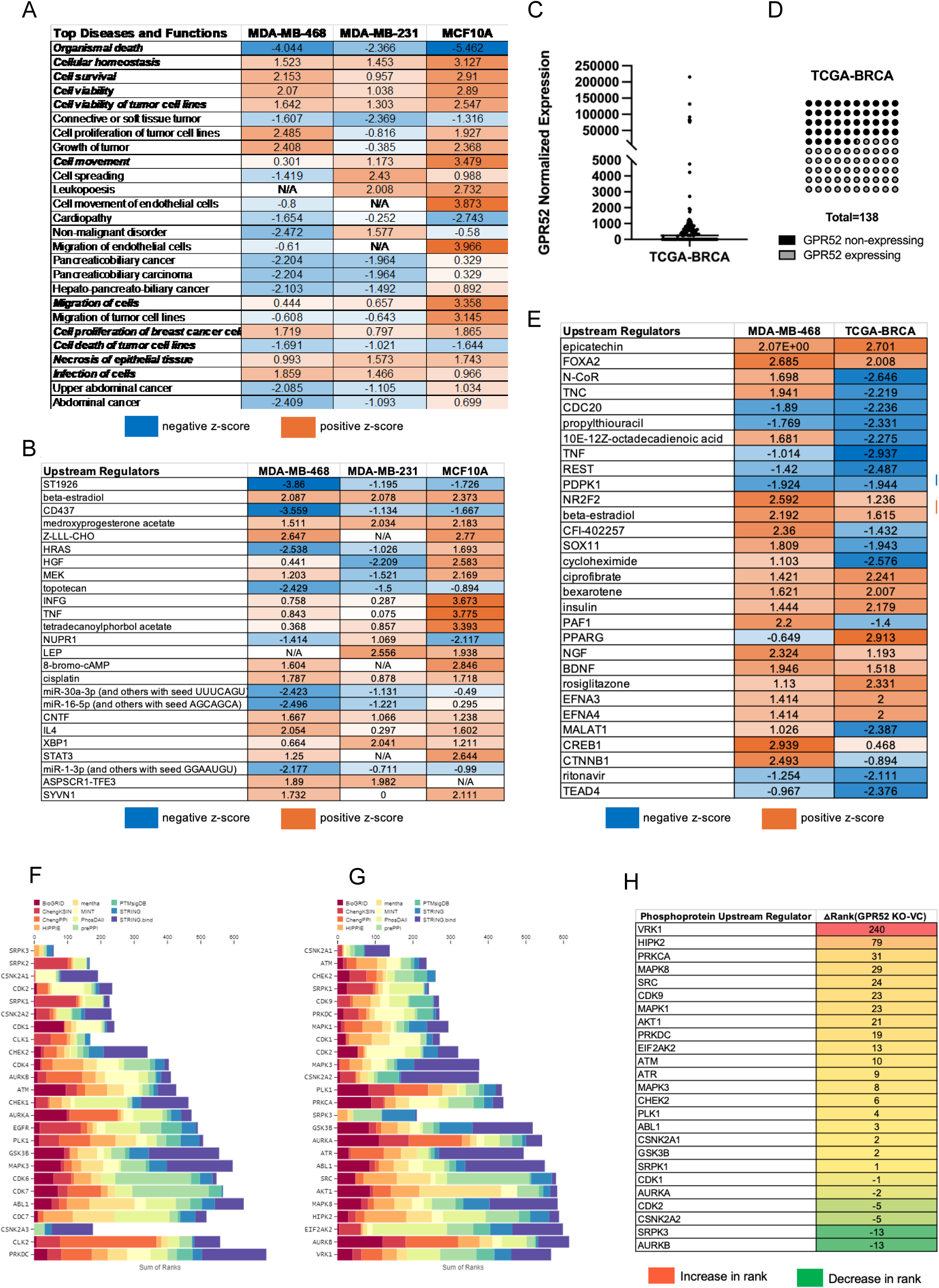
Proteomic and transcriptomic analyses of GPR52-null breast cell lines and resected breast tumors. (A-B) Differentially expressed proteins in the GPR52 KO cell lines with P-value <0.05, FDR<0.05, and fold-change greater than two were imported into IPA for each of the three cell lines. The common predicted upstream regulators (A) and top diseases and functions (B) with a P-value <0.05 are depicted. Pathways are marked N/A (not-applicable) if the z-score is not determined. (C) Normalized expression of GPR52 in TCGA-BRCA cohort resected breast tumors D) Graphical representation of proportion of patients with undetected (black) and detected (grey) GPR52 mRNA in resected breast tumors. E) DEGs in GPR52 KO MDA-MB-468 cells and the GPR52-null TCGA-BRCA cohort with P-value <0.05, FDR<0.05, and fold-change greater than two were imported into IPA for pathway analysis. Upstream regulators predicted to be responsible for the DEGs observed in both datasets with a P-value <0.05 are depicted. (F-H) Phosphoproteomic analysis was conducted on the groups outlined in Fig 5A. KEA3 software was used to rank kinases that were predicted to be active in the vector control (F) and GPR52 KO (G) cell lines. H) The difference in rank of kinases which were reported for both the GPR52 KO and vector control groups.

The TCGA-BRCA dataset reports RNA sequencing data of resected breast tumors from a cohort of patients. Normalized GPR52 mRNA expression in TCGA-BRCA tumors was visualized (Fig. 4C), and it was determined that 45.7% did not express GPR52, whereas the remainder expressed non-zero levels of GPR52 (Fig. 4D). The RNA-sequencing datasets were compared for the GPR52 non-expressing and -expressing cohorts and the differentially expressed genes (DEGs) were imported into IPA. We also conducted RNA sequencing of GPR52 KO and WT MDA-MB-468 cells that were cultured in a monolayer and imported the DEGs into IPA. We identified several common signatures in both datasets. Notably, upregulation of the signature associated with cAMP response element-binding protein (CREB1), a transcription factor that is activated downstream of cAMP signaling, was increased in both datasets (Fig. 4E) (19). Based on phosphoproteomic analyses of MDA-MB-468, MDA-MB-231, and MCF10A cells, we identified kinases predicted to be active in the vector control (Fig. 4F) and GPR52 KO (Fig. 4G) datasets (20). The rank of each of the top kinases in the GPR52 KO cell lines was then compared to that in the vector control group (Fig. 4H). This revealed the greatest increase in the rank of VRK1, which regulates cell cycle progression via phosphorylation and activation of CREB (21).

### cAMP production is regulated by GPR52 and mediates phenotypes associated with GPR52 loss

Next, we explored whether cAMP could modulate the phenotypes observed following GPR52 loss in MDA-MB-468 cells. GPR52 KO cells exhibited less rounding and were able to spread on plastic with adenylyl cyclase inhibitor (ACi) treatment (Fig. 5A; arrows). Interestingly, the expression of Snai1 was induced by forskolin (FSK, a direct activator of adenylyl cyclase) treatment of WT cells, whereas vimentin expression was reduced in GPR52 KO cells treated with ACi (Fig. 5B). ACi treatment of GPR52 KO MDA-MB-231 cells also increased the area of DQ normalized to the area of tdTomato without affecting the degree of colocalization of DQ and tdTomato (Fig. 5C-E).

**Figure 5.**
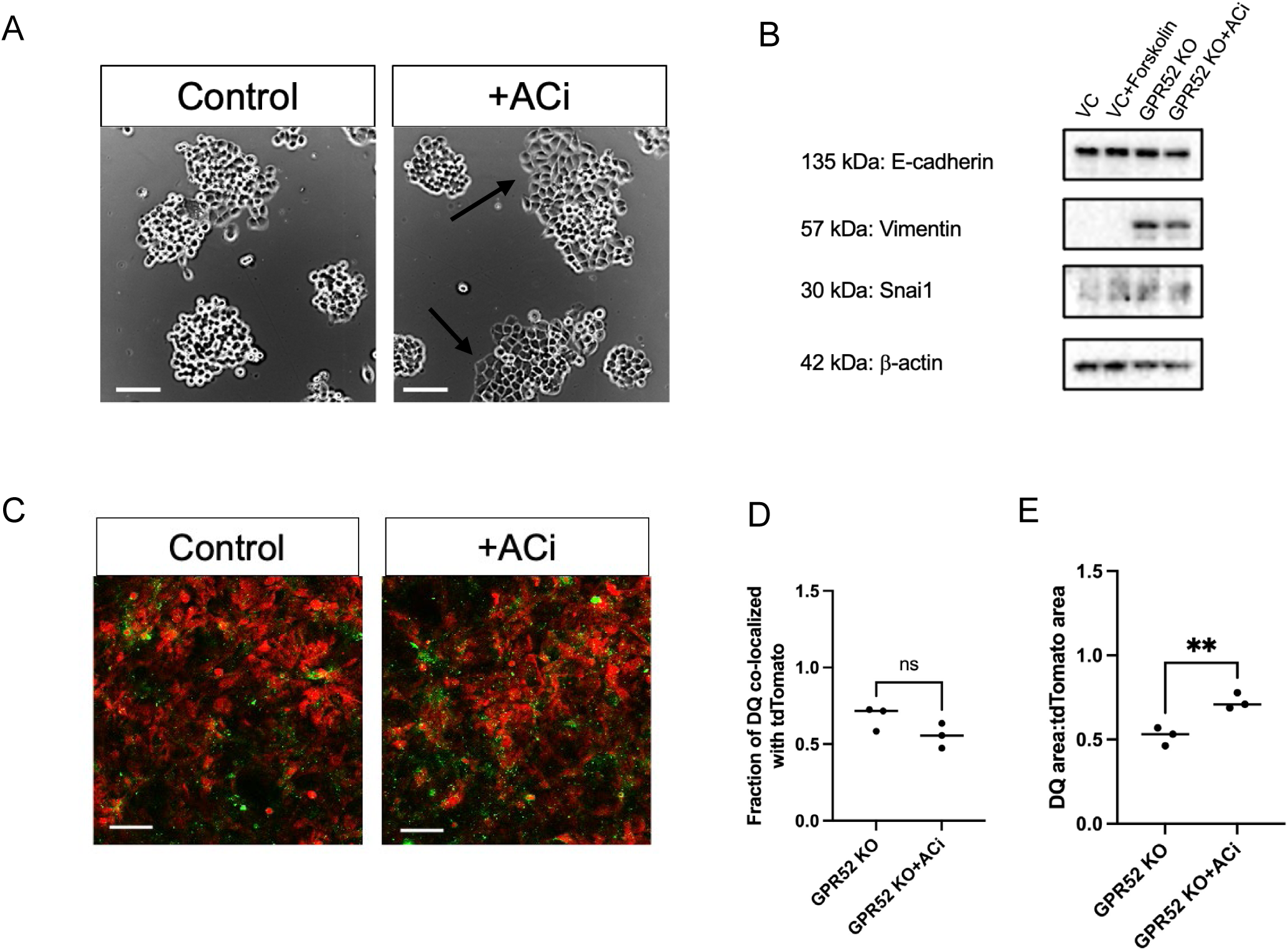
cAMP partially mediates phenotypes associated with GPR52 loss. A) GPR52 KO (sgRNA2) MDA-MB-468 cells were cultured in monolayer and treated with 1.4 μM ACi or the vehicle control. Treatment was replaced every 72 hours over 6-8 days. Scalebar=100 mm. B) Western blotting of MDA-MB-468 cells treated for 24 hours. C) GPR52 KO MDA-MB-231 cells were plated to confluency in monolayer and treated the next day with 1.4 mM ACi or vehicle control. Invasion assays were conducted as in Fig. 3. Scalebar=100 mm. The fraction of DQ co-localized with tdTomato (D) and area of DQ normalized to the area of tdTomato (E) at z=10 mm at t=24 hours. n=3, Student’s t-test, P-value<0.05; \**P* <0.05, \*\**P* <0.005, ns=not significant. Line=median.

To further explore the relationship between GPR52 and cAMP signaling, we next determined whether GPR52 activation led to a change in intracellular cAMP levels, as described by other groups (8, 9). To do this, we used a bioluminescence resonance energy transfer 2 (BRET2)-based EPAC sensor, GFP10-EPAC-RlucII (Supplementary Fig. S2A) (22). This modified form of EPAC contains a luminescent donor and a fluorescent acceptor that are in proximity to one another when EPAC is not bound to cAMP. However, the binding of cAMP induces a conformational shift that leads to an increased distance between the donor and acceptor, and a reduction in energy transfer. We found that increasing the amount of GPR52 and/or the concentration of FTBMT, a synthetic GPR52 agonist, led to a reduction in the BRET ratio in HEK293 cells (Supplementary Fig. S2B). Next, we introduced FTBMT and the EPAC sensor into MDA-MB-468 cells and found that BRET values tended to increase at higher doses (Supplementary Fig. S2C). To investigate whether GPR52 couples with Gaq, we also quantified intracellular Ca^2+^ levels over a similar range of FTBMT doses using the Ca^2+^-sensitive dye Fluo-4. Despite observing a robust response to the sarcoendoplasmic reticulum calcium ATPase (SERCA) inhibitor thapsigargin, we observed no change in baseline or thapsigargin-induced cytoplasmic Ca^2+^ levels (Supplementary Fig. S2D).

### Expression of the melanoma cell adhesion molecule (MCAM) is inversely related to GPR52 in breast cancer and is regulated by cAMP

RNA-sequencing analysis of MDA-MB-468 cells demonstrated that GPR52 loss is associated with differential expression of many cell adhesion molecules (CAMs) (Fig. 6A), including melanoma cell adhesion molecule (MCAM). MCAM is highly expressed in large blood vessels, but recent studies have also described increased MCAM expression in certain cancer types and its promotion of cancer progression (23–25). In breast cancer, increased MCAM mRNA expression in resected tumors was associated with a reduction in overall survival (Fig. 6B) (12). The expression of MCAM and GPR52 mRNA was inversely correlated in resected breast tumors (Fig. 6C). We found that MCAM protein expression was increased in GPR52 KO MDA-MB-468 cells and that 1.4 mM ACi treatment reduced MCAM expression (Fig. 6D). This was consistent with the effects of GPR52 loss in MCF10A cells (Supplementary Fig. S3).

**Figure 6.**
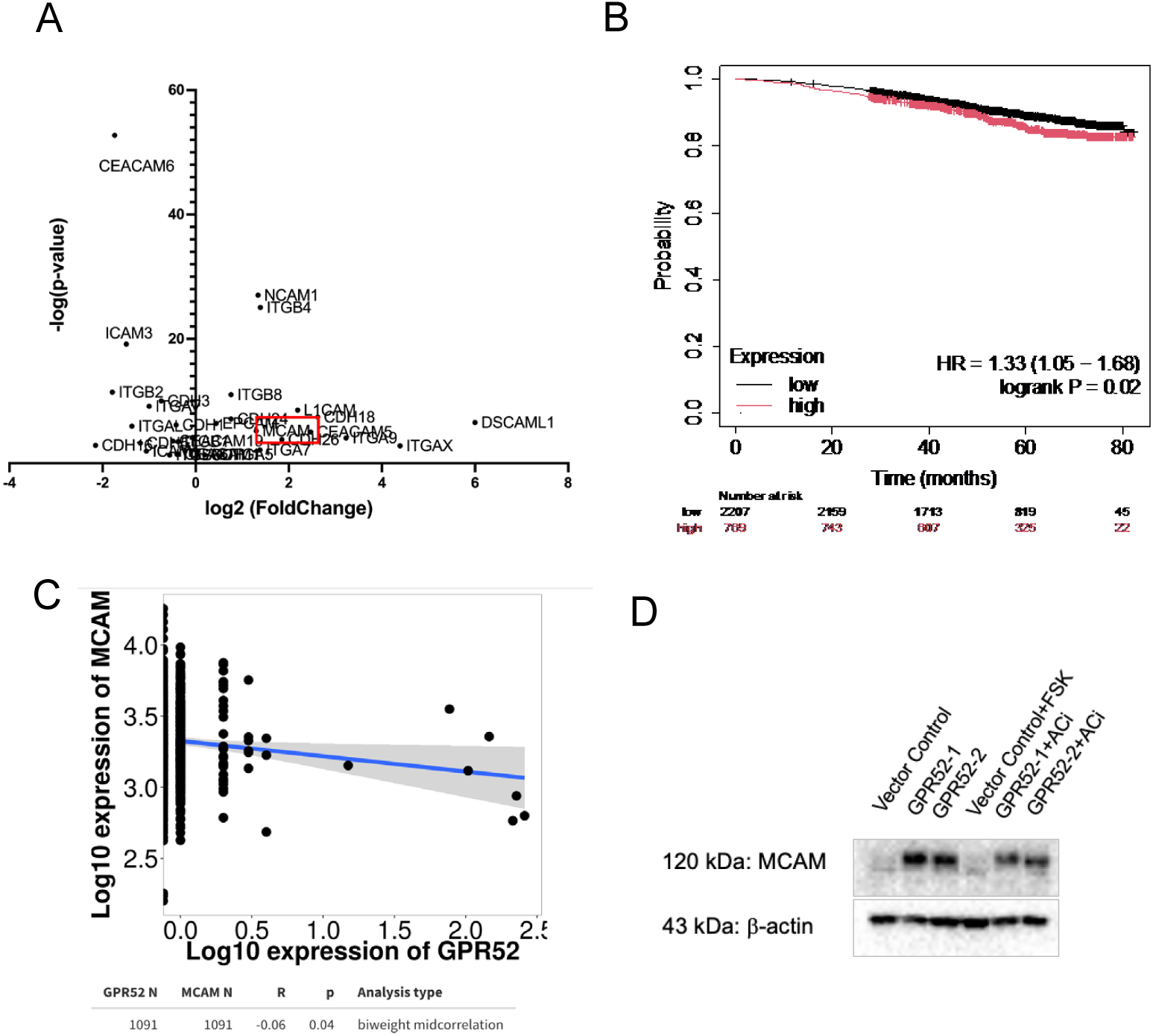
Expression of the melanoma cell adhesion molecule (MCAM) is inversely related to GPR52 in breast cancer and is regulated by cAMP. (A) CAMs that were differentially expressed in GPR52 KO MDA-MB-468 cells based on RNA-sequencing. P-value<0.05. n=3. (B) KMplot breast cancer RNA-seq webtool overall survival curves for patients with low versus high MCAM mRNA expression in resected tumors for all breast cancer subtypes. Low versus high cutoff was determined based on the maximum segregation between the groups. (C) TNMplot correlation of MCAM and GPR52 transcript expression in breast tumors from an RNA-sequencing dataset. (D) MDA-MB-468 cells were cultured in monolayer (TC-treated, non-PDL coated) and treated with forskolin (FSK), SQ22536 (ACi), or vehicle control for 24 hours. Cell lysates were probed for MCAM and beta-actin.

### Loss of GPR52 is associated with increased TNBC burden in zebrafish

As we observed that loss of GPR52 led to changes in organization, cell-cell adhesion, and invasiveness of breast cancer cells, we wanted to determine whether these changes were associated with an increase in breast cancer burden *in vivo.* To enable close monitoring of cancer cell organization and distribution, we used a zebrafish xenograft model (Fig. 7A) (26, 27). The TG(flk1:EGFP-NLS) zebrafish strain, which constitutively expresses a green fluorescent protein in endothelial cells, allows cancer cells to be monitored in relation to the zebrafish vasculature (28).

**Figure 7.**
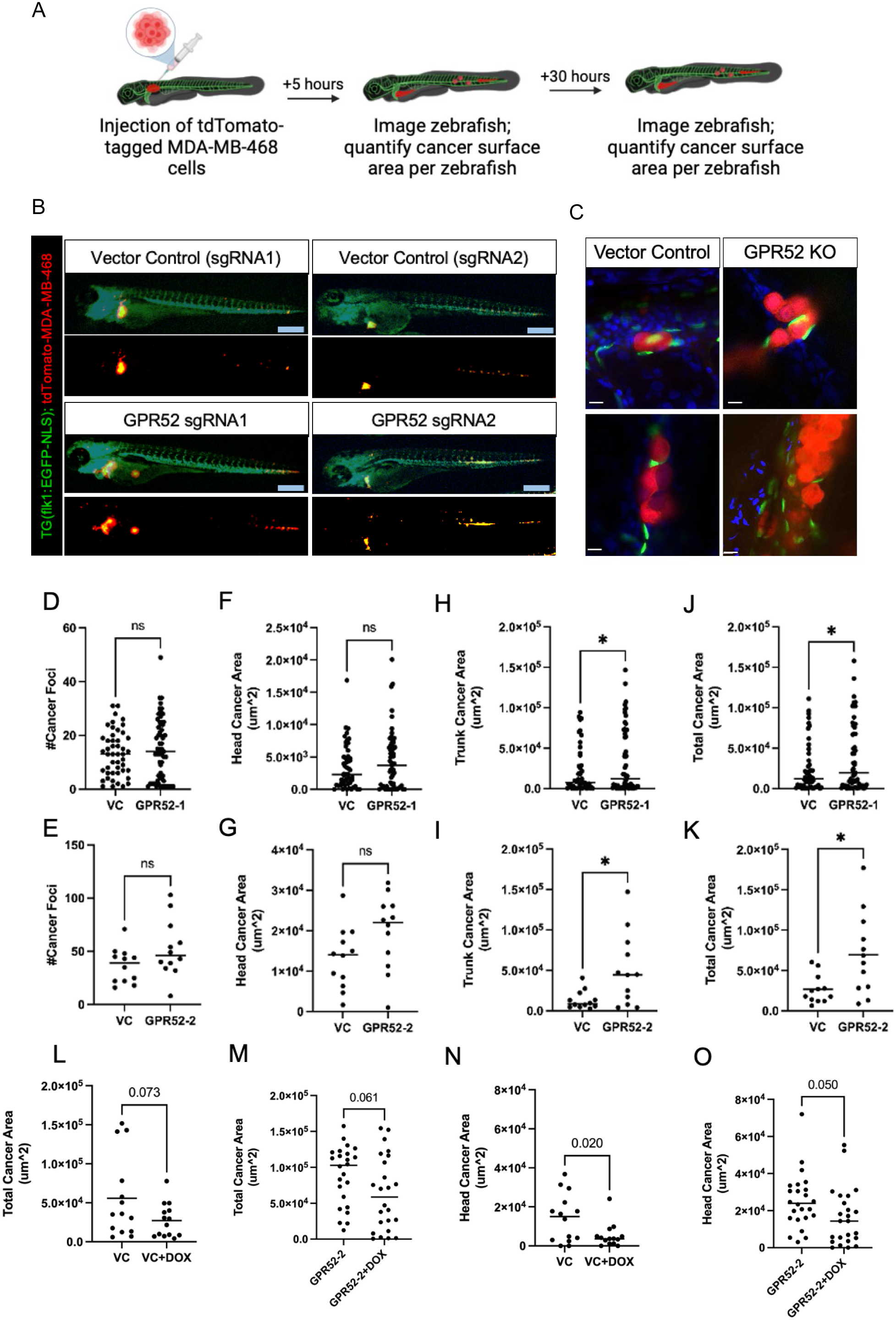
Loss of GPR52 is associated with increased TNBC burden in zebrafish. A) Schematic of zebrafish xenograft study. B) Tg(flk1:EGFP-NLS) zebrafish at 30 hpi. MDA-MB-468 cells (red) and endothelial cells (green) are visualized. Scalebar=300 μm. C) Visualization of the interaction between MDA-MB-468 cells (red) and endothelial cells (green) by confocal microscopy. Scalebar=10 μm. (D-E) The total number of tdTomato foci per zebrafish. (F-G) The total area of tdTomato in the head and (H-I) trunk per zebrafish. (J-K) The sum of the area of tdTomato in the head and trunk is expressed as the total cancer area per zebrafish. VC=vector control. n=51 VC-1, n=59 GPR52-1, n=12 VC-2, n=12 GPR52-2. Student’s t-test, P<0.05. *= *P*<0.05. ns=not significant. Line=median. (L-M) Zebrafish were treated with 8 mM doxorubicin or vehicle control (VC) and imaged at 30 hpi. The total area of tdTomato per zebrafish and (N-O) the area of tdTomato in the zebrafish head were determined. VC=vector control. DOX=doxorubicin. n=14 VC, n=24 GPR52-2. Student’s t-test, P<0.05. ns=not significant. Line=median.

Two independent studies were performed to compare the behavior of the vector control MDA-MB-468 cells with GPR52 sgRNA1 and sgRNA2 cells. We found that WT and GPR52 KO MDA-MB-468 cells were detectable throughout the body of the zebrafish at 30 h post-injection (hpi) (Fig. 7B). We also observed that WT and GPR52 KO cells circulated collectively in the zebrafish bloodstream and identified endothelial cells between cancer cells, suggesting an interaction between the two cell types (Fig. 7C). The number of cancer foci did not differ significantly between groups at 30 hpi (Fig. 7D-E). However, the total cancer area, which was calculated as the sum of the area of cancer in the head (superior to the otolith) and trunk (distal to the injection site, not including the yolk sac) (Fig. 7F-I),) was significantly greater in zebrafish injected with GPR52 KO cells (Fig. 7J-K).

Increased clustering of cancer cells is associated with reduced sensitivity to cytotoxic chemotherapeutic drugs, particularly when used as a single agent (2, 29). Doxorubicin is one of the most potent chemotherapeutic drugs approved by the Food and Drug Administration and is used in breast cancer treatment (30, 31). Therefore, we designed a zebrafish xenograft therapeutic study that incorporated 8 μM doxorubicin or the vehicle control (Milli-Q water) in E3 water maintained at 5 hpi and then quantified the cancer area at 30 hpi (32). At 30 hpi, there was a trend towards reduced total cancer area in doxorubicin-treated animals for both WT and GPR52 KO groups (Fig. 7L-M). Doxorubicin caused a significant reduction in the breast cancer cell area in the head of the zebrafish for both WT and GPR52 KO cells, with noticeable residual disease in animals xenografted with GPR52 KO cells (Fig. 7N-O).

## Discussion

The data presented herein demonstrate that GPR52 is a novel regulator of multicellular organization in breast cancer and its loss can promote features of cancer progression. First, we observed an increase in cell-cell interactions and cell clustering in 2D cultures with GPR52 loss in MCF10A, MDA-MB-468, and MDA-MB-231 cell lines. TEM demonstrated that GPR52 loss in MDA-MB-468 cells is associated with a reduction in the length of the interface between the two cells, but that GPR52 KO cells are more juxtaposed with one another than WT cells, which exhibit more intercellular space along their interface. Multicellular aggregation and increased cell-cell adhesion are mechanisms by which cancer cells can increase their metastatic potential (2). Moreover, the increase in disorganization and number of MDA-MB-468 spheroids with GPR52 loss are two hallmarks of cancer progression, with the latter suggesting a potential increase in stemness due to an increased propensity to survive and proliferate in Matrigel (33).

Furthermore, collective organization and invasion provide mechanisms for cancer cells to transmit survival signals and invade directionally, in some cases featuring a distinct leading front of cells that tends to be more mesenchymal and invasive (34). We also found that GPR52 loss affected the pattern and extent to which cancer cells. A computational model has previously suggested that collective invasion requires less ECM digestion than single-cell invasion (35). Linearization and alignment of collagen tracts have been hypothesized to promote the directed migration of cancer cells, and the alignment of ECM fibers has been associated with a reduction in proteolytic degradation of the ECM (35, 36).

Many upstream regulators of EMT have been identified; however, to our knowledge, induction of EMT or partial EMT with the loss of a GPCR has not been previously described. Importantly, partial EMT has been documented in circulating tumor cells (37). The exclusivity of our observations of cell elongation and the development of mesenchymal morphology on poly-D-lysine, but not tissue culture-treated plastic, suggests that integrin activity can influence the extent of EMT in this cell line and is an interesting area for further exploration.

Our finding that GPR52 loss was associated with an increase in the total cancer area in zebrafish at 30 hpi suggests increased survival and/or growth of GPR52 KO MDA-MB-468 cells. Our data also suggested a reduction in the sensitivity of cancer cells in the head to doxorubicin with GPR52 loss. Notably, the blood-brain barrier starts to develop 3 days post-fertilization (dpf) in zebrafish (38). As doxorubicin has a limited capacity to cross the blood-brain barrier, its efficacy has historically been limited to its effects outside the CNS (39). However, blood-brain barrier-permeable forms of doxorubicin have been developed and may have notable utility against head or brain metastases for certain molecular subtypes of breast cancer, as demonstrated in this study (39).

We identified an increased cAMP signaling signature in GPR52 KO groups based on RNA sequencing and proteomic studies and found that modulation of cAMP levels could induce or attenuate some, but not all, phenotypes associated with GPR52 loss. Increased intracellular cAMP levels have previously been associated with increased proliferation in 3D cultures, increased invasiveness of breast cancer, and deposition of ECM components (13, 40). However, the role of cAMP in regulating ECM degradation has not been previously characterized. Therefore, these mechanistic studies not only provide some insight into how GPR52 loss may influence breast cancer cell biology but also identify the role of cAMP in regulating breast cancer cell adhesion and ECM digestion. Treatment of HEK293 cells with FTBMT led to an increase in intracellular cAMP, as previously reported, but MDA-MB-468 cells demonstrated a lack of change or possible decrease in intracellular cAMP with FTBMT treatment and no change in calcium levels (41). The coupling of GPCRs to G proteins is promiscuous, and one study estimated that 73% of GPCRs can activate multiple G proteins (42). Therefore, the interaction between GPR52 and different G proteins, particularly in different cell types, warrants further investigation.

Furthermore, we found that the melanoma cell adhesion molecule (MCAM), which is considered a potential biomarker and promoter of the progression of many cancers, was increased in MCF10A and MDA-MB-468 cells, and that its expression could be reduced in GPR52 KO cells with adenylyl cyclase inhibitor treatment (43, 44). Thus, we identified a mechanism by which MCAM is upregulated in breast cancer and a method to reduce the expression of this driver of cancer aggression.

### Conclusion

Our study identified GPR52 as a regulator of cancer cell clustering, collective organization, invasion, and EMT. Using a zebrafish xenograft model, we demonstrated that loss of GPR52 is associated with increased tumor burden. RNA sequencing and proteomic analyses of WT and GPR52-null breast cancer cells demonstrated differences in many gene and protein signatures, including increased cAMP signaling in GPR52-null groups. We showed that inhibition of cAMP production can rescue some phenotypes associated with GPR52 loss. Thus, we provide a rationale for investigating the therapeutic effect of GPR52 agonism in breast cancer and encourage investigation of the role of GPR52 in the progression of additional cancer types. Furthermore, our work identifies novel features of cell biology that can be regulated by GPCR and broadens the scope of significance of this class of proteins in cancer biology.

## List of abbreviations

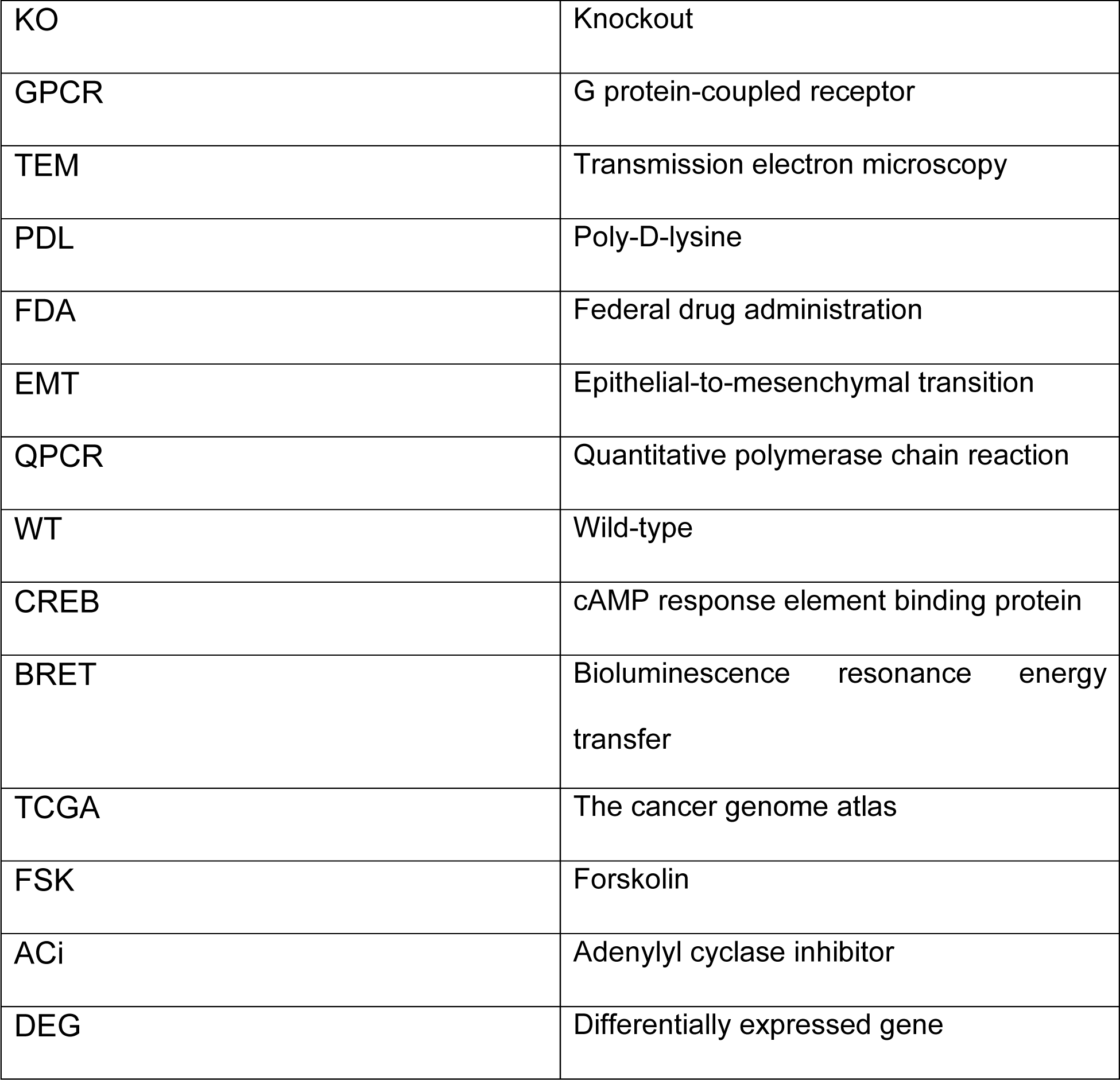
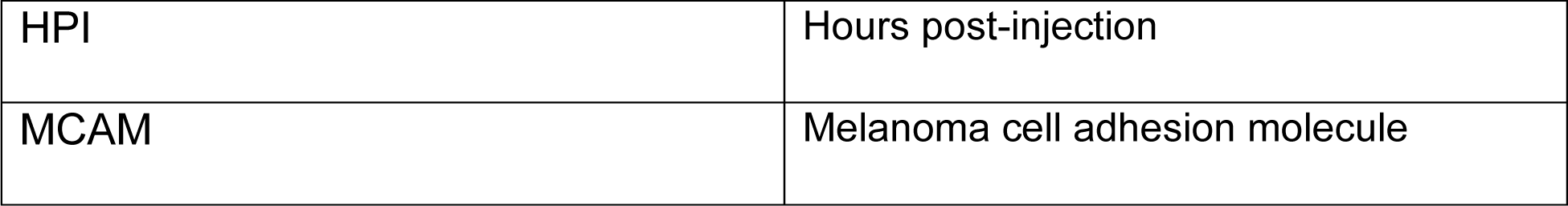

## Declarations

### Ethics approval and consent to participate

All animal experiments were performed with the approval of the Institutional Animal Care and Use Committee (IACUC) protocol #2011-0100 from Weill Cornell Medicine with animal care under the supervision of the Research Animal Resource Center. Approval date: September 19, 2023.

### Consent for publication

Not applicable

### Availability of data and materials

The datasets used and/or analyzed during the current study are available from the corresponding author upon reasonable request. Proteomics and phoshphopeptide-enriched data were submitted to the MassIVE Proteomics Repository. Data are available via MassIVE (MSV000065698, MSV000095699) or ProteomeXchange (PXD055208, PXD055209) for global and phosphopeptide-enriched data, respectively.

## Author’s Contributions

SZH conceptualized, designed, and wrote the manuscript with KAB. SZH performed the majority of the experiments in this study, except for those specified hereafter. CCA has developed CRISPR-Cas9 tools for GPR52 expression. IT performed the cancer cell injections in zebrafish. SYJ and TF performed RNA-seq analyses for the TCGA datasets. BB and OE performed RNA-seq analysis for MDA-MB-468 WT vs. GPR52 KO cells. MW performed proteomic analyses. DD and THE conducted the BRET studies. SL assisted with the tissue culture. PB optimized and provided the protocols for MCF10A culture. TE-supervised zebrafish studies. PKA, RT, and SP conducted calcium fluo-4 experiments. LD provided the CRISPR plasmids, tools, and study advice. JB provided intellectual advice and materials for conducting this research.

## Acknowledgements

Studies were conducted with the support and facilities provided by the Microscopy and Image Analysis Core Facility and the Genomics Resources Core Facility at Weill Cornell Medicine, and the Mass Spectrometry and Proteomics Core Facility at Kansas University Medical Center. All schematics were created using BioRender.

## Materials and Methods

### Cell culture

Human breast cell lines MDA-MB-468 (RRID:CVCL_0419), MDA-MB-231 (RRID:CVCL_0062), MCF7 (RRID:CVCL_0031), HS578T (CVCL_0332), T47D (CVCL_0553), and MCF10A (RRID:CVCL_0598) were purchased from the American Type Culture Collection (ATCC). MDA-MB-468, MDA-MB-231, MCF7, and HS578T cells were cultured in Dulbecco’s Modified Eagle’s medium (DMEM, Gibco #11995-065) supplemented with 10% fetal bovine serum (FBS) (Gibco 26140079, Sigma-Aldrich F0926) and 1% penicillin-streptomycin ( complete medium). T47D cells were cultured in Roswell Park Memorial Institute medium (RPMI 1640, Thermo Fisher Scientific #11875) supplemented with 10% fetal bovine serum (FBS) (Gibco 26140079) and 1% penicillin-streptomycin ( complete medium). MCF10A cells were cultured in Dulbecco’s Modified Eagle’s Medium/F12 (Gibco, #11330-032) supplemented with 5% FBS, 1% penicillin-streptomycin, and the following growth factors: epidermal growth factor (20 ng/ml), hydrocortisone (0.5 mg/ml), cholera toxin (100 ng/ml), and insulin (10 μg/ml) (i.e., complete media). All cell lines were maintained at 37°C in a humidified atmosphere containing 5% CO2. All cell lines were authenticated by STR analysis and tested regularly for mycoplasma contamination using the Universal Mycoplasma Detection Kit (ATCC 30-1012K) and MycoAlert Mycoplasma Detection Kit (Lonza LT07-318).

### Breast cancer survival curves

The KM plotter RNA-seq breast cancer web tool (www.kmplot.com) (12) was used to obtain Kaplan-Meier survival curves. The associated hazard ratios, 95% confidence intervals, and log-rank P-values were automatically determined using the web tool. Low versus high cut-off values were determined based on the maximum segregation between the groups.

### GPR52 and MCAM expression comparison in cancerous and non-cancerous human tissue

GPR52 mRNA expression in cancerous and non-cancerous tissues was analyzed across solid cancer types using RNA-seq datasets from the TNMplot database and a web tool (https://www.tnmplot.com) (11). The following datasets were analyzed for this analysis: adrenocortical carcinoma (“adrenal”), bladder urothelial carcinoma (“bladder”), breast invasive carcinoma (“breast”), colon adenocarcinoma (“colon”), esophageal carcinoma (“esophageal”), kidney renal clear cell carcinoma (“renal_cc”), kidney renal papillary cell carcinoma (“renal_pa”), liver hepatocellular carcinoma (“liver”), lung adenocarcinoma (“lung_ac”), lung squamous cell carcinoma (“lung_sc”), ovarian serous cystadenocarcinoma (“ovary”), pancreatic adenocarcinoma (“pancrease”), prostate adenocarcinoma (“prostate”), rectum adenocarcinoma (“rectum”), skin cutaneous melanoma (“skin”), stomach adenocarcinoma (“stomach”), testicular germ cell adenocarcinoma (“testis”), thyroid carcinoma (“thyroid”), uterus corpus endometrial carcinoma (“uterus_ec”). GPR52 expression between normal, tumor, and metastatic breast-derived samples was compared based on a gene chip dataset available through the TNM plot. Source data for GPR52 expression were collected in January 2024, and the correlation between GPR52 and MCAM expression in breast tumors was determined in June 2024.

### RT-qPCR

Each breast cancer cell line was cultured in an individual well of a 6-well tissue culture-treated plate until 50-80% confluent. Total RNA was extracted using an RNeasy Mini Kit (Qiagen # 74106), and RNA purity was assessed using a Nanodrop 2000 instrument (Thermo Fisher Scientific). RNA was reverse-transcribed using the qScript cDNA Synthesis kit (VWR # 101414-100) based to the manufacturer’s protocol. Quantitative polymerase chain reaction (PCR) was performed in triplicate from the RNA collected from each well using Fast SYBR Green Master Mix (Thermo Fisher Scientific #4385612) following the standard protocol. GPR52 and RPL32 transcripts were quantified using the following primers.

GPR52 F: 5’ - CGGGTCTTGGACAATCCAACTC - 3’,

GPR52 R: 5’ - TGCTTCCTGATCCTTCACACAC - 3’,

RPL32 F: 5’ – CAGGGTTCGTAGAAGATTCA - 3’,

RPL32 R: 5’ – CTTGGAGGAAACATTGTGAGCGATC - 3’.

GPR52 expression in each human breast cell line was normalized to the housekeeping gene RPL32, and then normalized to the average expression of GPR52 across the human breast cell lines analyzed.

### Generation of stable cell lines

Transduction-based CRISPR-Cas9 gene editing was used to generate indels in *GPR52* in MDA-MB-468, MDA-MB-231, and MCF10A cells. For lentivirus production, HEK293 cells were plated in a six-well plate and transfected with a prepared mix in 150uL DMEM (with no supplements) containing 2.5 μg of pLenti-Cas9-P2A-Puro (a gift from Lukas Dow; RRID:Addgene_110837), 1.25 μg of PAX2, 1.25 μg of VSV-G, and 30 μl of polyethylenimine (1 mg/ml). The medium was replaced the following day and changed 36 h post-transfection to the target cell collection medium. MDA-MB-468, MDA-MB-231, and MCF10A cells were plated in individual wells of a six-well plate and transduced with the pLenti-Cas9-P2A-Puro lentivirus generated above in serum- and antibiotic-free media containing 8 mg/mL polybrene for 24 h, and then selected with 1-2.5 mg/mL puromycin in complete media (45).

The forward and reverse sgRNA cloning primers were as follows:

sgRNA1-F: CACCGCAAAACCATGGCGTAGCGA

sgRNA1-R: AAACTCGCTACGCCATGGTTTTGC

sgRNA2-F: CACCGTGAATGGTGTGCCACGTCT

sgRNA2-R: AAACAGACGTGGCACACCATTCAC

The primers were annealed and cloned following standard procedures using the BsmBI/EcoRI site of the pLenti-U6-sgRNA-tdTomato-P2A-BlasR (LRT2B) lentiviral vector (a gift from Lukas Dow; RRID:Addgene_110854) (45). MDA-MB-468, MDA-MB-231, and MCF10A cells stably expressing the pLenti-Cas9-P2A-Puro lentiviral vector were transduced with GPR52 sgRNA lentiviral vectors in serum- and antibiotic-free media containing 8 mg/mL polybrene for 24 h and subjected to blasticidin selection followed by limiting dilution assays to isolate single cells from the sgRNA-transduced populations. The cells were then expanded to generate clonal populations, and a subset of these cells was collected by centrifugation for DNA extraction and Sanger sequencing of the GPR52 sgRNA target region. Briefly, DNA was extracted from the cell pellet using the QiaAMP DNA Mini Kit (Qiagen #56304) according to the manufacturer’s protocol. The GPR52 guide target regions were then amplified by PCR using the following primers and purified using the QIAquick PCR Purification Kit (Qiagen #28104).

GPR52-1 F: 5’ - AAACCTGGTTACCATGGTGAC - 3’,

GPR52-2 F: 5’ - GATCTGGATCTACTCCTGCC - 3’,

GPR52-1,-2 R: 5’ - ATTATATAGGGGAGCCACAGC-3’,

The amplified PCR products were subjected to Sanger sequencing (Genewiz, Azenta) to identify clones with indels in GPR52 sgRNA target regions (Supplementary Fig. S1). The clonal populations generated from each sgRNA were expanded and used in downstream studies. A heterogeneous population of gSafe cells was used as the vector control population.

### Transmission electron microscopy

MDA-MB-468 cells were cultured in suspension in non-tissue culture-treated plates for 24-48 hours and then briefly centrifuged in 1.7mL tubes. The resulting pellet was washed and fixed with 4% paraformaldehyde, 2.5% glutaraldehyde, and 0.02% picric acid in 0.1 M sodium cacodylate buffer (pH 7.3). The samples were stained with uranyl acetate and dehydrated using a graded ethanol series. After dehydration, the samples were covered with a layer of fresh resin, and embedded molds were inserted into the resin. After polymerization, the samples were cut at 200 nm for screening by light microscopy and then at 65 nm to be mounted on grids for TEM under a JEOL JSM 1400 (JEOL, USA) electron microscope. The camera used was a Veleta 2 K × 2 K CCD (EMSIS, GmbH, Muenster). Images were taken at 2500-20,000× magnification, and each field included the extracellular space, cytoplasmic area, and nucleus. Cell-cell interface length was determined using 2500x images by drawing a straight line that extended the distance of the cell-cell interface only between cells that exhibited extensive adhesion between their plasma membrane surfaces. The cell diameter was determined using 2500x images by drawing a straight line that spanned the midpoint of the aforementioned analyzed cells. The free intercellular space was determined using Fiji software. The image threshold intensity was set to distinguish the cells from the (white) background. The boundary between the two cells was then outlined, and the fraction of the area occupied by free space (met the threshold intensity) was determined.

### 3D Matrigel cell culture and analysis

MDA-MB-468 cells were resuspended in 1:1 growth-factor-reduced Matrigel:serum-free DMEM and plated at a density of 10,000 cells/100 μL per well in a black optical-bottom 96-well plate (Corning Matrigel #356231). Complete medium (100 μL) was added to each well after solidification of the Matrigel, and the medium was replaced every 2-3 days. After 10 days of culture, the cells were fixed with 10% formalin and stained with Hoechst 33342 (Santa Cruz Biotechnology # SC-495790). Cells were imaged using a confocal microscope (Zeiss LSM880) to detect cytoplasmic tdTomato and nuclear Hoechst signals. Images were then imported to Imaris Microscopy Image Analysis Software (OXFORD instruments). The Imaris “Surface” function was used to determine the area per spheroid, total area occupied per well, and roundness of each spheroid based on the absolute intensity of the tdTomato signal. For MDA-MB-468 cells, the following formula for circularity was used to determine the roundness of each spheroid.

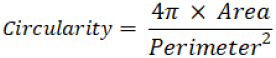

### Matrigel invasion assays

MDA-MB-468 and MDA-MB-231 cells were plated at >80% confluence in optical grade 96-well plates. A 50 μL solution of 1:1 growth-factor-reduced Matrigel:serum-free DMEM ± 1:40 DQ-Collagen IV (Thermo Fisher Scientific) was added above the monolayer and allowed to form a gel. Fifty microliters of DMEM containing 10% FBS was added above the gel once it solidified. After 24 h, confocal microscopy (Zeiss LSM880) was used to obtain a z-stack of tdTomato and DQ signals in each well at 10 μm intervals starting from the base of the well. The Imaris “Surface” function was used to determine the surface area of tdTomato foci above the minimum fluorescence intensity threshold at the established z-planes. For collective invasion analyses and class B determination at z=10um, the threshold intensity was auto-determined by Imaris per image. Surfaces were identified based on the local contrast of the tdTomato signal with the non-fluorescent background. The Imaris “Surface” function was used to quantify the area of DQ at z=10 μm based on a maintained absolute value threshold across samples and colocalize the volume of DQ with the volume of tdTomato. that was adjusted to maximize detection across samples based on the local contrast at z=10 μm.

### Poly-D-lysine coating

PDL (Santa Cruz Biotechnology #SC-136156) was resuspended in Milli-Q water at 1mg/mL and frozen in aliquots at −20°C. To develop coating solutions, PDL was resuspended in Milli-Q water to a final concentration of 100 μg/mL and added to tissue culture-treated plates to cover the surface. plates were then incubated for at least 30 min at room temperature or several hours at 4°C. The plates were washed twice with PBS and used for cell culture experiments.

### Western blotting and analysis

Cells were cultured in monolayers under standard tissue culture conditions as described above. At the time of collection, cells were scraped on ice with ice-cold PBS and centrifuged at 1500-2000 rpm, and the resulting cell pellet was snap-frozen in liquid nitrogen. On the day of western blotting, the cell pellet was lysed in ice-cold buffer (5 mM HEPES, 137 mmol/L NaCl, 1 mmol/L MgCl2, 1 mmol/L CaCl2, 10 mmol/L NaF, 2 mmol/L EDTA, 10 mmol/L sodium pyrophosphate, 2 mmol/L NaVO4, 1% NP-40, 10% glycerol) containing protease inhibitors (Thermo Fisher Scientific #78429), incubated on a rotating shaker at 4°C for 15 min, and centrifuged for 20 min at 4°C at 20,000 rpm. Cell extracts were denatured in buffer containing β-mercaptoethanol, run on NuPAGE 4–12% Bis-Tris protein gels (Thermo Fisher Scientific), and then transferred to nitrocellulose membranes. Membranes were blocked with 5% nonfat dry milk (Bio-Rad, #1706404) for one hour at room temperature (RT), washed with 1X Tris-Buffered Saline, 0.1% Tween (TBST), and incubated with primary antibodies at 4°C for at least 16 h. The following primary antibodies were used: E-cadherin (RRID:AB_2291471), Snai1 (RRID:AB_2255011), vimentin (RRID:AB_10695459), p-CREB (RRID:AB_2561044), CREB (RRID:AB_310268) and MCAM (RRID:AB_2143373). The membranes were then washed with TBST and incubated with secondary antibodies for 1-2 hours at room temperature. The secondary antibodies used were anti-rabbit IgG HRP-linked antibody (RRID:AB_2099233) and anti-mouse IgG HRP-linked antibody (RRID:AB_330924). Western Lightning Plus ECL (Thermo Fisher Scientific #509049325) and ImageLab software (Bio-Rad) were used for band detection. Membranes were stripped with Restore PLUS Western blot stripping buffer (Thermo Fisher Scientific, #46430) for 15 min and then probed for β-actin expression as a loading control using an HRP-linked β-actin antibody (RRID:AB_262011).

### Zebrafish maintenance

All animal experiments were performed under an approved IACUC protocol (2011–0100) from Weill Cornell Medicine, with animal care under the supervision of the Research Animal Resource Center. The TG(flk1:EGFP-NLS) zebrafish strain was generated by the Markus Affolter Laboratory (Biozentrum, University of Basel, Switzerland) (Developmental Biology 316:312-322, 2008) and kindly provided by Jesus Torres-Vazquez (New York University, New York, NY). Embryos were obtained by natural matings and raised at 28.5 °C with a 14-h light–10-h dark cycle. Zebrafish were maintained in E3 water (5 mM NaCl, 0.17 mM KCl, 0.33 mM CaCl2, 0.33 mM MgSO4, 1 ppm methylene blue).

### Zebrafish xenograft studies

tdTomato-tagged MDA-MB-468 cells were passed through a 40 μm filter and resuspended in PBS at a density of 400 cells/5nL. Two dpf zebrafish embryos were each injected with 5nL of cancer cells in the perivitelline space or the duct of Cuvier, whichever was best accessible based on the orientation of the fish. Injected embryos were evaluated at two hpi and only zebrafish with detectable tdTomato distal to the injection site were maintained at 32 °C. Zebrafish were anesthetized with 0.02% tricaine and imaged at five hpi and 30 hpi using a Nikon SMZ1500 microscope mounted with a Nikon DS-FI3 camera. Cancer surface area per zebrafish was quantified using Imaris “Surface” function; two rectangles were drawn per zebrafish to encompass the area distal to the injection site excluding the yolk sac (i.e., trunk) and superior to the otolith (i.e., head), and the total surface area of tdTomato determined in each rectangle was combined to calculate the total cancer area. The number of detected tdTomato surfaces was used to determine the number of cancer foci. Zebrafish were fixed with 4% paraformaldehyde at the study endpoint. For confocal imaging, fixed zebrafish were washed with PBS and then incubated for at least 12 h at 4°C in PBS containing Hoechst 33342. The zebrafish were then mounted in 1% low-melt agarose and imaged using a confocal microscope (Zeiss LSM880).

For drug treatment studies, zebrafish with detectable tdTomato distal to the yolk sac were selected at two hpi, placed into individual wells of 24-well plates, and imaged at 5 hpi. The zebrafish were then maintained in E3 water containing either 8 μM doxorubicin or a vehicle (Milli-Q water). All zebrafish were imaged at 30 hpi and analyzed as described above.

### RNA-sequencing studies and computational analyses

MDA-MB-468 WT (parental) and GPR52 sgRNA1 KO cells (one mixed population and two clonal populations) were cultured in 6-well plates. Total RNA was extracted using the QIAzol lysis reagent (Qiagen #79306) and RNeasy Mini Kit (Qiagen #74106). Samples were subjected to RNA sequencing at the Genomics Resources Core Facility (GRCF, Weill Cornell Medicine). Total RNA integrity was assessed with a 2100 Bioanalyzer (Agilent Technologies, Santa Clara, CA). RNA concentration was measured using a NanoDrop system (Thermo Fisher Scientific, Inc., Waltham, MA, USA). RNA sample library and RNA sequencing were performed by the Genomics Core Laboratory at Weill Cornell Medicine with the Illumina TruSeq Stranded mRNA Sample Library Preparation kit (Illumina, San Diego, CA), according to the manufacturer’s instructions. The normalized cDNA libraries were pooled and sequenced on an Illumina NovaSeq 6000 sequencer with 50 paired-end cycles. The raw sequencing reads in BCL format were processed using bcl2fastq 2.19 (Illumina) for FASTQ conversion and demultiplexing.

All reads were independently aligned with STAR_2.4.0f1 for sequence alignment against the human genome sequence build hg19 downloaded using the UCSC genome browser and SAMTOOLS v0.1.19 for sorting and indexing reads. Cufflinks (2.0.2) was used to estimate the expression values (FPKMS) and GENCODE v19 GTF files for annotation. Gene counts from the htseq-count and DESeq2 Bioconductor packages were used to identify DEGs. All DEGs with P-value <0.05 and false discovery rate <0.05 were uploaded to Ingenuity Pathway Analysis software (Qiagen) for further analysis.

### Proteomic studies and computational analyses

Parental, vector control, and GPR52-1 and −2 KO MDA-MB-468, MDA-MB-231, and MCF10A cells were cultured in 10-cm tissue-culture-treated plates until 60-80% confluent. Cells were then scraped on ice with ice-cold PBS and centrifuged, and the resulting cell pellet was snap-frozen in liquid nitrogen. Cells were lysed by resuspension in radioimmunoprecipitation assay (RIPA) buffer supplemented with protease and phosphatase inhibitors. Soluble proteins were reduced by the addition of TCEP (0.5 M) to a final concentration of 5mM followed by incubation at 55 °C for 30 min. The reduced samples were alkylated by the addition of 375 mM iodoacetamide to a final concentration of 10 mM, followed by incubation in the dark at room temperature for 30 min. Ice-cold acetone was added to each sample at a volume ratio of 5:1. The samples were vortexed and stored at −20 °C overnight. After precipitation, the samples were centrifuged at 14,000 × *g* and 4 °C for 30 min to pellet the proteins. The supernatant was removed, and the pellet was air-dried on a benchtop for 10 min. Proteins were resuspended in 50 mM TEAB (pH 8) containing 2 mM CaCl2. Trypsin was added (500 ng), and the proteins were allowed to digest overnight at 37 °C with shaking at 500 RPM (Thermomixer, Eppendorf). Digestion was quenched by adding 10% formic acid to a final concentration of 1%. Digested samples were centrifuged at 10,000 × *g* for 10 min to remove particulates, and the supernatant was transferred to a fresh tube and stored at −20 °C until phosphopeptide enrichment.

The samples were enriched for phosphorylated peptides using SMOAC. Briefly, digested samples were first enriched using the High Select™ Phosphopeptide Enrichment Kit (Thermo Fisher Scientific) following the manufacturer’s protocol. The flow-through was applied to the High Select™ Fe-NTA Phosphopeptide Enrichment Kit (Thermo Fisher Scientific), following the manufacturer’s protocol. After the second enrichment, the flow-through became the global, unenriched sample, and the elutes from both kits were pooled to generate the phosphopeptide-enriched fraction. The peptide concentration was measured at 205 nm using a NanoDrop spectrophotometer (Thermo Scientific).

Samples were injected using the Vanquish Neo (Thermo Fisher Scientific) nano-UPLC onto a C18 trap column (0.3 mm x 5 mm, 5 µm C18) using pressure loading. Peptides were eluted onto a separation column (PepMap™ Neo, 75 µm × 150 mm, 2 µm C18 particle size, Thermo Fisher Scientific) prior to elution directly onto the mass spectrometer. Briefly, peptides were loaded and washed for 5 min at a flow rate of 0.350 µL/min in 2% B (mobile phase A: 0.1% formic acid in water; mobile phase B: 80% ACN, 0.1% formic acid in water). Peptides were eluted over 100 minutes from 2-25% mobile phase B before ramping to 40% B for 20 min. The column was washed for 15 min at 100% B before reequilibration at 2% B for the next injection. The nano-LCs were directly interfaced with an Orbitrap Ascend Tribrid mass spectrometer (Thermo Fisher Scientific) using a silica emitter (20 µm i.d., 10 cm) equipped with a high-field asymmetric ion mobility spectrometry (FAIMS) source. Data were collected by data-dependent acquisition with the intact peptide detected in the Orbitrap at 120,000 resolving power from to 375-1500 *m/z*. Peptides with charge +2-7 were selected for fragmentation by higher-energy collision dissociation (HCD) at 28% NCE and were detected in the ion trap using a rapid scan rate (global) or Orbitrap at a resolving power of 30,000 (enriched). Dynamic exclusion was set to 60s after one instance. The mass list is shared among the FAIMS compensation voltages. FAIMS voltages were set at −45 (1.4 s), −60 (1 s), −75 (0.6 s) CV for a total duty cycle time of 3s. Source ionization was set at 1700 V with an ion transfer tube temperature of 305 °C. Raw files were searched against the human protein database downloaded from UniProt on 05-05-2023 using SEQUEST in Proteome Discoverer 3.0. The abundances, abundance ratios, and P-values were exported to Microsoft Excel for further analysis. All proteins with differential GPR52 KO vs. WT abundance ratios (P-value <0.05, false discovery rate <0.05) were uploaded to the Ingenuity Pathway Analysis software (Qiagen) for further analysis of each of the three cell lines. KEA3 software was used to rank the kinase activity of the combined vector control and GPR52 KO populations across the three cell lines based on their respective phosphoproteomes (20).

### Analysis of TCGA-BRCA transcriptomic dataset

The Cancer Genome Atlas (TCGA)-BRCA database was used to obtain primary breast tumor gene expression data from 142 patients with breast cancer. The downloaded data displayed the gene ENSEMBL ID and FPKM-UQ normalized expression counts. Four patients were found to have mutations in the *Gpr52* gene and were not included in the downstream analyses. The data for the remaining 138 patients were segregated based on “zero” and “non-zero” expression of GPR52 and then subsequently read into edgeR, a software package available from BiocManager for differential expression analysis of RNA-sequencing data (46). Briefly, dispersion was estimated to measure the inter-library variability. The data were then fitted to a generalized linear regression model. Statistical significance testing (likelihood ratio test) between the cohorts was performed using the fitted model to compare differential gene expression between “zero” and “non-zero data,” creating the dataset “Non-zero/Zero”. All DEGs with P-value <0.05 and false discovery rate <0.05 were uploaded to Ingenuity Pathway Analysis software (Qiagen) for further analysis.

### BRET biosensor studies

HEK 293 and MDA-MB-468 cells were maintained in DMEM with 10% FBS, respectively, and cultured at 37°C and 5% CO2. For transfection, cells were plated at a density of 2.5 × 10^5^ cells (HEK293) and 8 × 10^5^ cells (MDA-MB-468) per well in 6-well plates (Thermo Scientific, 140,675). On the day of transfection, the medium was replaced with DMEM containing 2.5% FBS (HEK 293) and Opti-MEM (MDA-MB-468) (Thermo Fisher Scientific #31985062) without antibiotics. HEK293 cells were transfected using Lipofectamine 2000 (Thermo Fisher Scientific, #11668027). The GFP10-Epac-RlucII in pcDNA3.1 (generously provided by Dr. Michel Bouvier, Université de Montréal) and GPR52 cDNA ORF Clone GPR52 (Sino Biological # HG25891-UT) in pcDNA3.1, with added N-terminal flag tag, were used in these studies.

For HEK293 cells, per well: 1.5 μg of plasmid DNA (total) was added to 100 μl DMEM (no serum, no antibiotics) in a 1.5 mL Eppendorf tube. In another tube, 3 μl of Lipofectamine 2000 was added to 100 μl of DMEM (no serum or antibiotics).

For MDA-MB-468 cells, per well: 2.5 μg of plasmid DNA (total) was added to 100 μl Opti-MEM (no serum, no antibiotics) in a 1.5 mL Eppendorf tube. In another tube, 5 μl of Lipofectamine 2000 or 3000 was added to 100 μl of Opti-MEM (no serum or antibiotics).

After 5 min, the contents of the tubes were combined and gently mixed or vortexed for each cell line. The mixture was incubated for 20-30 minutes at room temperature. The 200 μL DNA:Lipofectamine mixture was then added dropwise per well, and the plate was swirled gently and incubated at 37 °C and 5% CO2 for 4-5 hours. The medium was then replaced with a complete medium.

Cells were detached the day following transfection with 0.25% trypsin–EDTA (Wisent) and plated at a density of 3 × 10^4^ cells/well in a poly-l-ornithine (Sigma-Aldrich)-coated 96-well white bottom plate (Thermo Scientific, 236,105) for BRET analysis. After 24 h of incubation in the 96-well plate, the media was removed and cells were washed once with Krebs buffer (146 mM NaCl, 42 mM MgCl2, 10 mM HEPES pH 7.4, 1 g/L D-glucose) and incubated for 2 h at room temperature in Krebs buffer. Coelenterazine 400A (Cedarlane) was added, to a final concentration of 2.5 μM, to each well, followed by incubation for 5 min prior to the basal reading. FTBMT was added to the indicated final concentration and the plate was read after 15 or 30 min of stimulation. MDA-MB-468 cells were co-stimulated with 1 mM forskolin.

The BRET ratios were calculated as the emission at 515 nm and 410 nm. For all experiments, ΔBRET = (BRET ratio from cells stimulated with FTBMT and forskolin) – (BRET ratio from vehicle treated with forskolin). Three technical replicates were used for all the treatments. The BRET experiments were performed using a Tristar2 plate reader (Berthold. Technologies GmbH and Co. KG). The normalized BRET ratio was computed as BRETStimuated/BRETVehicle. Dose-response curves were plotted using non-linear regression. Data are presented as the mean ± standard error (SE). For MDA-MB-468 cells, the experiment was conducted five times, and the aggregate data from all experiments are presented. For HEK 293 cells, the experiment was conducted twice, and the combined data from the two experiments are presented.

### Intracellular calcium measurements

MDA-MB-468 cells were seeded at a density of 5000 cells/well in a 96-well plate. After 24 h, the cell-permeable calcium indicator Fluo-4 (Thermo Fisher Scientific, #F14201) was added to each well. cells were incubated for 45 min at 37 °C, then for 15 min at room temperature. Baseline fluorescence measurements were obtained (t=0), and FTBMT was added to select the wells. The plate was read at 30 second intervals for 10 min. Thapsigargin (Sigma-Aldrich) was added to cells as a positive control, and the plate was then read at 30 second intervals for an additional 10 min (47).

### Drug preparation

Forskolin (Santa Cruz Biotechnology #SC-3562) and SQ22536 (Santa Cruz Biotechnology #SC-201572) were resuspended in ethanol (1 mM) and DMSO (14 mM), and frozen. SQ22536 was aliquoted to avoid freeze-thaw cycles. Ethanol and DMSO were used as vehicle controls in all experiments that incorporated these compounds.

### Statistical analysis

All data were expressed as individual data points with a line at the median value or as the mean ± SEM, unless indicated otherwise. The groups were compared using statistical tests for significance, as described in the figure legends. Statistical significance was set at P <0.05. Statistical tests were performed using the GraphPad Prism software.

The data generated in this study are available upon request from the corresponding author.

## Supplemental figures

**Supplementary Figure S1.**
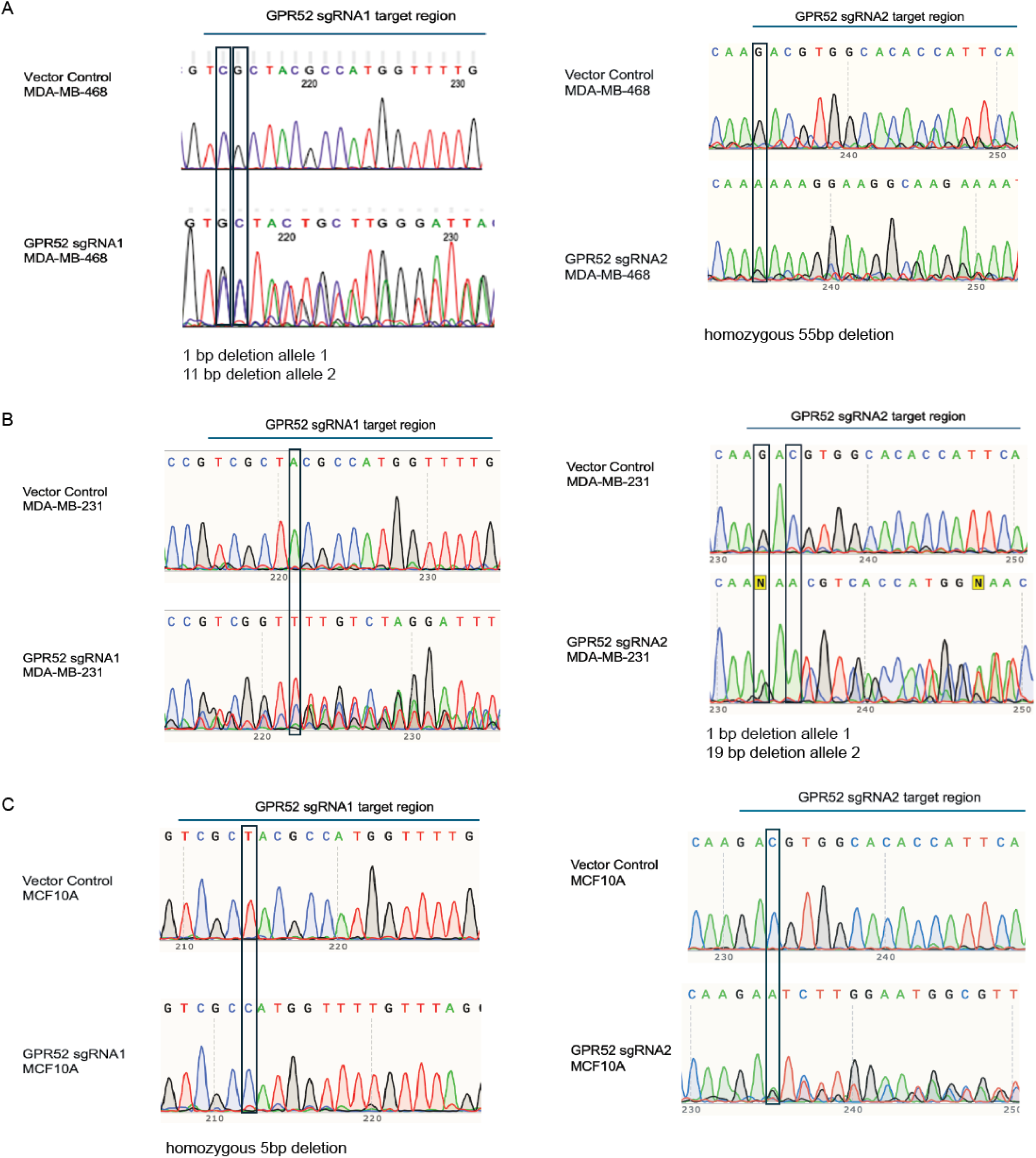
Sanger sequencing of breast cell lines to confirm indels in the *GPR52* gene. MDA-MB-468 (A), MDA-MB-231 (B), and MCF10A (C) were transduced with Cas9 followed by a GPR52-targeted sgRNA or the empty vector. DNA was extracted from cell pellets and sequenced using GPR52 primers described in *Materials and Methods.* Indels in the sgRNA target regions were confirmed for cell lines transduced with GPR52 sgRNA1 and sgRNA2, and the nucleotide position indicating loss of the WT allele is outlined in black.

**Supplementary Figure S2.**
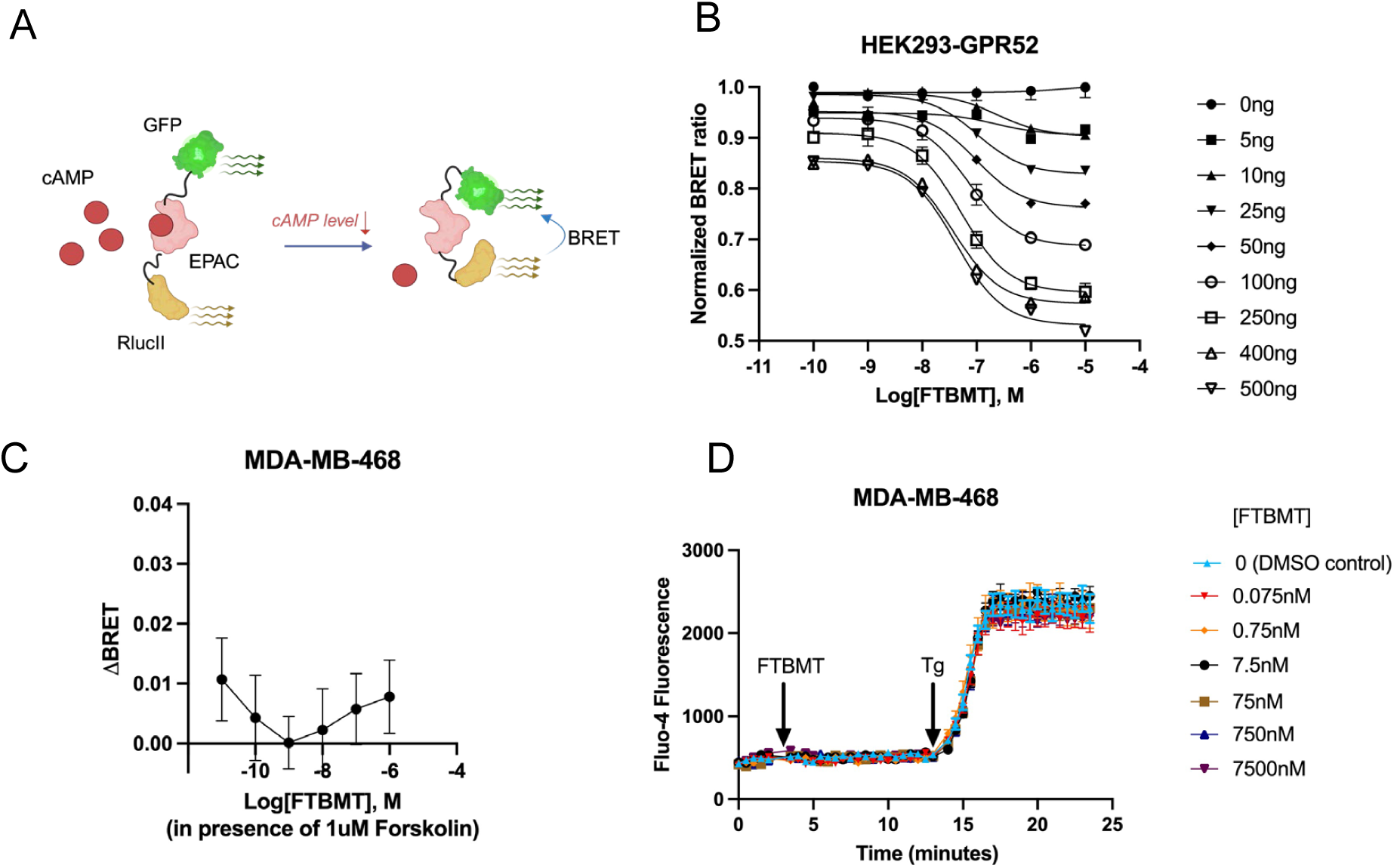
Treatment with GPR52 agonist FTBMT in HEK293 and MDA-MB-468 cells. A) Schematic of BRET-based GFP10-EPAC1-RlucII cAMP sensor. B) HEK293 cells transfected with GPR52 and GFP-EPAC1-RlucII were treated with varying concentrations of FTBMT for 15 minutes. Datapoints=mean, error bars=SEM. Data is pooled from 1-2 experiments, n=2 technical replicates per experiment. C) MDA-MB-468 cells transfected with the EPAC1 sensor were treated with increasing concentrations of FTBMT for 30 minutes in the presence of 1 μM forskolin to assess for possible G⍺i coupling. Datapoints=mean, error bars=SEM. Data was pooled from five independent experiments, n=3 technical replicates per experiment. D) MDA-MB-468 cells were plated at a density of 5,000 cells/well in a 96-well plate. After 24 hours, the cell-permeable calcium indicator Fluo-4 was added to each well. Baseline fluorescence measurements were then obtained (t=0) and FTBMT was added at varying concentrations to achieve the final concentrations listed. The plate was read at 30 second intervals for 10 minutes. 100 nM Thapsigargin (Tg) was then added to all wells as a positive control for [Ca^2+^] detection. The plate was then read for 30 second intervals for an additional 10 minutes. Datapoints=mean, error bars=SEM. n=8.

**Supplementary Figure S3.**
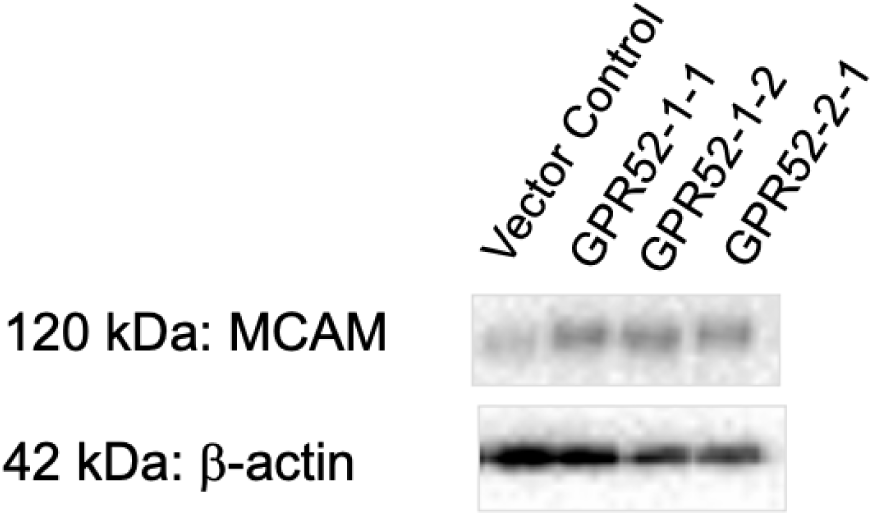
Detection of MCAM in MCF10A cell lysate. MCF10A vector control, two unique isogeneic GPR52 sgRNA1-transduced cell lines, and GPR52-2 cells were cultured in monolayer to 50-85% confluency. Cell lysates were collected as described in *Materials and Methods* and probed for MCAM and beta-actin.

## References

1. Hagemeister FB, Jr., Buzdar AU, Luna MA, Blumenschein GR. Causes of death in breast cancer: a clinicopathologic study. Cancer. 1980;46(1):162–7.

2. Schuster E, Taftaf R, Reduzzi C, Albert MK, Romero-Calvo I, Liu H. Better together: circulating tumor cell clustering in metastatic cancer. Trends Cancer. 2021;7(11):1020–32.

3. Bakir B, Chiarella AM, Pitarresi JR, Rustgi AK. EMT, MET, Plasticity, and Tumor Metastasis. Trends Cell Biol. 2020;30(10):764–76.

4. Venter JC, Adams MD, Myers EW, Li PW, Mural RJ, Sutton GG, et al. The sequence of the human genome. Science. 2001;291(5507):1304-51.

5. Hanson MA, Stevens RC. Discovery of new GPCR biology: one receptor structure at a time. Structure. 2009;17(1):8–14.

6. Hamm HE, Gilchrist A. Heterotrimeric G proteins. Curr Opin Cell Biol. 1996;8(2):189–96.

7. Chaudhary PK, Kim S. An Insight into GPCR and G-Proteins as Cancer Drivers. Cells. 2021;10(12).

8. Komatsu H, Maruyama M, Yao S, Shinohara T, Sakuma K, Imaichi S, et al. Anatomical transcriptome of G protein-coupled receptors leads to the identification of a novel therapeutic candidate GPR52 for psychiatric disorders. PLoS One. 2014;9(2):e90134.

9. Nishiyama K, Suzuki H, Harasawa T, Suzuki N, Kurimoto E, Kawai T, et al. FTBMT, a Novel and Selective GPR52 Agonist, Demonstrates Antipsychotic-Like and Procognitive Effects in Rodents, Revealing a Potential Therapeutic Agent for Schizophrenia. J Pharmacol Exp Ther. 2017;363(2):253–64.

10. Yao Y, Cui X, Al-Ramahi I, Sun X, Li B, Hou J, et al. A striatal-enriched intronic GPCR modulates huntingtin levels and toxicity. Elife. 2015;4.

11. Bartha A, Gyorffy B. TNMplot.com: A Web Tool for the Comparison of Gene Expression in Normal, Tumor and Metastatic Tissues. Int J Mol Sci. 2021;22(5).

12. Gyorffy B. Survival analysis across the entire transcriptome identifies biomarkers with the highest prognostic power in breast cancer. Comput Struct Biotechnol J. 2021;19:4101–9.

13. Au CC, Furness JB, Britt K, Oshchepkova S, Ladumor H, Soo KY, et al. Three-dimensional growth of breast cancer cells potentiates the anti-tumor effects of unacylated ghrelin and AZP-531. Elife. 2020;9.

14. Sriram K, Insel PA. G Protein-Coupled Receptors as Targets for Approved Drugs: How Many Targets and How Many Drugs? Mol Pharmacol. 2018;93(4):251–8.

15. Hughes CS, Postovit LM, Lajoie GA. Matrigel: a complex protein mixture required for optimal growth of cell culture. Proteomics. 2010;10(9):1886–90.

16. Guan JL, Trevithick JE, Hynes RO. Fibronectin/integrin interaction induces tyrosine phosphorylation of a 120-kDa protein. Cell Regul. 1991;2(11):951–64.

17. Kornberg L, Earp HS, Parsons JT, Schaller M, Juliano RL. Cell adhesion or integrin clustering increases phosphorylation of a focal adhesion-associated tyrosine kinase. J Biol Chem. 1992;267(33):23439–42.

18. Jedeszko C, Sameni M, Olive MB, Moin K, Sloane BF. Visualizing protease activity in living cells: from two dimensions to four dimensions. Curr Protoc Cell Biol. 2008;Chapter 4:Unit 4 20.

19. Gonzalez GA, Montminy MR. Cyclic AMP stimulates somatostatin gene transcription by phosphorylation of CREB at serine 133. Cell. 1989;59(4):675–80.

20. Kuleshov MV, Xie Z, London ABK, Yang J, Evangelista JE, Lachmann A, et al. KEA3: improved kinase enrichment analysis via data integration. Nucleic Acids Res. 2021;49(W1):W304–W16.

21. Kang TH, Park DY, Kim W, Kim KT. VRK1 phosphorylates CREB and mediates CCND1 expression. J Cell Sci. 2008;121(Pt 18):3035–41.

22. Martin RD, Sun Y, Bourque K, Audet N, Inoue A, Tanny JC, et al. Receptor-and cellular compartment-specific activation of the cAMP/PKA pathway by alpha(1)-adrenergic and ETA endothelin receptors. Cell Signal. 2018;44:43–50.

23. An Y, Wei N, Cheng X, Li Y, Liu H, Wang J, et al. MCAM abnormal expression and clinical outcome associations are highly cancer dependent as revealed through pan-cancer analysis. Brief Bioinform. 2020;21(2):709–18.

24. Wang J, Tang X, Weng W, Qiao Y, Lin J, Liu W, et al. The membrane protein melanoma cell adhesion molecule (MCAM) is a novel tumor marker that stimulates tumorigenesis in hepatocellular carcinoma. Oncogene. 2015;34(47):5781–95.

25. Zhang X, Wang Z, Kang Y, Li X, Ma X, Ma L. MCAM expression is associated with poor prognosis in non-small cell lung cancer. Clin Transl Oncol. 2014;16(2):178–83.

26. Teng Y, Xie X, Walker S, White DT, Mumm JS, Cowell JK. Evaluating human cancer cell metastasis in zebrafish. BMC Cancer. 2013;13:453.

27. Ren J, Liu S, Cui C, Ten Dijke P. Invasive Behavior of Human Breast Cancer Cells in Embryonic Zebrafish. J Vis Exp. 2017(122).

28. Blum Y, Belting HG, Ellertsdottir E, Herwig L, Luders F, Affolter M. Complex cell rearrangements during intersegmental vessel sprouting and vessel fusion in the zebrafish embryo. Dev Biol. 2008;316(2):312–22.

29. Campenni M, May AN, Boddy A, Harris V, Nedelcu AM. Agent-based modelling reveals strategies to reduce the fitness and metastatic potential of circulating tumour cell clusters. Evol Appl. 2020;13(7):1635–50.

30. Tacar O, Sriamornsak P, Dass CR. Doxorubicin: an update on anticancer molecular action, toxicity and novel drug delivery systems. J Pharm Pharmacol. 2013;65(2):157–70.

31. Chemotherapy for Breast Cancer [Internet]. American Cancer Society. 2021. Available from: https://www.cancer.org/cancer/types/breastcancer/treatment/chemotherapy-for-breast-cancer.html. Accessed July 2024.

32. Gopal U, Monroe JD, Marudamuthu AS, Begum S, Walters BJ, Stewart RA, et al. Development of a Triple-Negative Breast Cancer Leptomeningeal Disease Model in Zebrafish. Cells. 2023;12(7).

33. Qu Y, Han B, Yu Y, Yao W, Bose S, Karlan BY, et al. Evaluation of MCF10A as a Reliable Model for Normal Human Mammary Epithelial Cells. PLoS One. 2015;10(7):e0131285.

34. Vilchez Mercedes SA, Bocci F, Levine H, Onuchic JN, Jolly MK, Wong PK. Decoding leader cells in collective cancer invasion. Nat Rev Cancer. 2021;21(9):592–604.

35. Kumar S, Kapoor A, Desai S, Inamdar MM, Sen S. Proteolytic and non-proteolytic regulation of collective cell invasion: tuning by ECM density and organization. Sci Rep. 2016;6:19905.

36. Han W, Chen S, Yuan W, Fan Q, Tian J, Wang X, et al. Oriented collagen fibers direct tumor cell intravasation. Proc Natl Acad Sci U S A. 2016;113(40):11208–13.

37. Papadaki MA, Mala A, Merodoulaki AC, Vassilakopoulou M, Mavroudis D, Agelaki S. Investigating the Role of CTCs with Stem/EMT-like Features in Metastatic Breast Cancer Patients Treated with Eribulin Mesylate. Cancers (Basel). 2022;14(16).

38. Fleming A, Diekmann H, Goldsmith P. Functional characterisation of the maturation of the blood-brain barrier in larval zebrafish. PLoS One. 2013;8(10):e77548.

39. Rousselle C, Clair P, Lefauconnier JM, Kaczorek M, Scherrmann JM, Temsamani J. New advances in the transport of doxorubicin through the blood-brain barrier by a peptide vector-mediated strategy. Mol Pharmacol. 2000;57(4):679–86.

40. Insel PA, Murray F, Yokoyama U, Romano S, Yun H, Brown L, et al. cAMP and Epac in the regulation of tissue fibrosis. Br J Pharmacol. 2012;166(2):447–56.

41. Wang P, Felsing DE, Chen H, Stutz SJ, Murphy RE, Cunningham KA, et al. Discovery of Potent and Brain-Penetrant GPR52 Agonist that Suppresses Psychostimulant Behavior. J Med Chem. 2020;63(22):13951–72.

42. Masuho I, Kise R, Gainza P, Von Moo E, Li X, Tany R, et al. Rules and mechanisms governing G protein coupling selectivity of GPCRs. Cell Rep. 2023;42(10):113173.

43. Wu Z, Wu Z, Li J, Yang X, Wang Y, Yu Y, et al. MCAM is a novel metastasis marker and regulates spreading, apoptosis and invasion of ovarian cancer cells. Tumour Biol. 2012;33(5):1619–28.

44. Chen J, Dang Y, Feng W, Qiao C, Liu D, Zhang T, et al. SOX18 promotes gastric cancer metastasis through transactivating MCAM and CCL7. Oncogene. 2020;39(33):5536–52.

45. Zafra MP, Schatoff EM, Katti A, Foronda M, Breinig M, Schweitzer AY, et al. Optimized base editors enable efficient editing in cells, organoids and mice. Nat Biotechnol. 2018;36(9):888-93.

46. Huber W, Carey VJ, Gentleman R, Anders S, Carlson M, Carvalho BS, et al. Orchestrating high-throughput genomic analysis with Bioconductor. Nat Methods. 2015;12(2):115–21.

47. Wrennall JA, Ahmad S, Worthington EN, Wu T, Goriounova AS, Voeller AS, et al. A SPLUNC1 Peptidomimetic Inhibits Orai1 and Reduces Inflammation in a Murine Allergic Asthma Model. Am J Respir Cell Mol Biol. 2022;66(3):271–82.

